# Cardiovascular disease causes proinflammatory microvascular changes in the human right atrium

**DOI:** 10.1101/2021.06.23.449672

**Authors:** Suvi Linna-Kuosmanen, Eloi Schmauch, Kyriakitsa Galani, Johannes Ojanen, Carles A. Boix, Tiit Örd, Anu Toropainen, Prosanta K. Singha, Pierre R. Moreau, Kristiina Harju, Adriana Blazeski, Åsa Segerstolpe, Veikko Lahtinen, Lei Hou, Kai Kang, Elamaran Meibalan, Leandro Z. Agudelo, Hannu Kokki, Jari Halonen, Juho Jalkanen, Jarmo Gunn, Calum A. MacRae, Maija Hollmén, Juha Hartikainen, Minna U. Kaikkonen, Guillermo García-Cardeña, Pasi Tavi, Tuomas Kiviniemi, Manolis Kellis

## Abstract

Ischemic heart disease is globally the leading cause of death. It plays a central role in the electrical and structural remodeling of the right atrium, predisposing to arrhythmias, heart failure, and sudden death. Here, we provide the first dissection of the gene expression changes in the live right atrial tissue, using single-nuclei RNA-seq and spatial transcriptomics. We investigate matched samples of the tissue and pericardial fluid and reveal substantial differences in disease-associated gene expression in all cell types, leading to inflammatory microvascular dysfunction and changes in the tissue composition. Our study demonstrates the importance of creating high-resolution cellular maps and partitioning disease signals beyond epicardial coronary arteries and ischemic left ventricle to identify candidate mechanisms leading to more severe types of human cardiovascular disease.

**One-Sentence Summary:** Single-cell dissection of *ex vivo* heart biopsies and pericardial fluid in ischemic heart disease and heart failure

## Main Text

Since the first landmark study of the single-cell dissection of adult human hearts in 2020 (*1*), several papers describing the cellular composition of the heart (*2–4*) and its transcriptional changes in cardiovascular disease (*5–9*) have been published. Yet, the effects of cardiovascular disease on the right atrium have not been investigated, although the sinus node – the natural pacemaker of the heart – resides in the roof of the right atrium (*10*). Given its central role in the normal cardiac function, and the unknown mechanisms by which chronic cardiovascular diseases cause electrical and structural remodeling of the atrial myocardium, it is important to dissect the molecular biology of the right atrium in health and disease, to improve the understanding of how these common diseases predispose to problems, such as sinus node dysfunction and arrhythmias, and increase the risk of heart failure and sudden death (*11–13*).

Ischemic heart disease is the leading cause of death globally according to the World Health Organization. It is commonly described as epicardial obstructive atherosclerotic coronary artery disease, although according to recent studies, fewer than one in five patients with known or suspected ischemic heart disease have obstructive disease (*14–17*). Instead, coronary microvascular dysfunction has been suggested as a major underappreciated and understudied condition underlying major cardiovascular diseases, including ischemic disease and arrhythmias, and shown to associate with more advanced disease and worse disease outcomes (*17–22*). Deciphering the role microvasculature plays in the disease manifestation and progression in functionally central parts of the heart seems imperative. Nevertheless, earlier studies have exclusively focused on the structural and functional abnormalities of epicardial arteries in patients with coronary artery disease, for these arteries are easily visible, whereas coronary microvasculature can only be studied indirectly (*17*). Coronary microvascular aberrations mediate ischemia and cause symptoms both in patients with obstructed and unobstructed coronary arteries (*23, 24*), and therefore, its effects are unlikely to be limited to the left ventricle, which is often seen as the epicenter of myocardial ischemia and cardiovascular disease.

Here, we present a comprehensive single-cell atlas of the *ex vivo* right atrium in ischemic heart disease and heart failure. We use single-nuclei RNA-seq (snRNA-seq), spatial transcriptomics, and human primary cell models to dissect the right atrium in the presence and absence of the diseases, revealing inflammatory changes in the tissue in response to disease manifestation and progression. We follow the findings in matched samples of right atrial tissue and pericardial fluid, comparing changes in stable disease, acute myocardial infarction, and after recovery from the acute phase, exploring similarities, differences and potential interactions between the tissue and fluid cells. Our expression module-based dissection of the disease-associated genes highlights inflammatory modules in the vasculature and the pericardial fluid cells, suggesting a genetic component to the observed inflammatory changes in connection to microvascular dysfunction.

## Results

### The first expression map of the ex vivo right atrium

We obtained *ex vivo* cardiac tissue biopsies from the right atrial appendage of 10 control individuals, 15 patients with ischemic heart disease (IHD), 9 with myocardial infarction (MI), 11 with ischemic heart failure (IHF), and 3 with non-ischemic heart failure (NIHF, i.e., patients with no coronary artery disease) (**Fig. 1a**). We profiled the samples with snRNA-seq and processed the data utilizing our custom method for cardiac tissue (*25*), resulting in 296 682 nuclei that were grouped into 12 main cell types (**Fig. 1b-f**), using available marker genes (*1–3*) and gene ontology enrichments of biological processes (**Fig. S1a**).

**Fig. 1.**
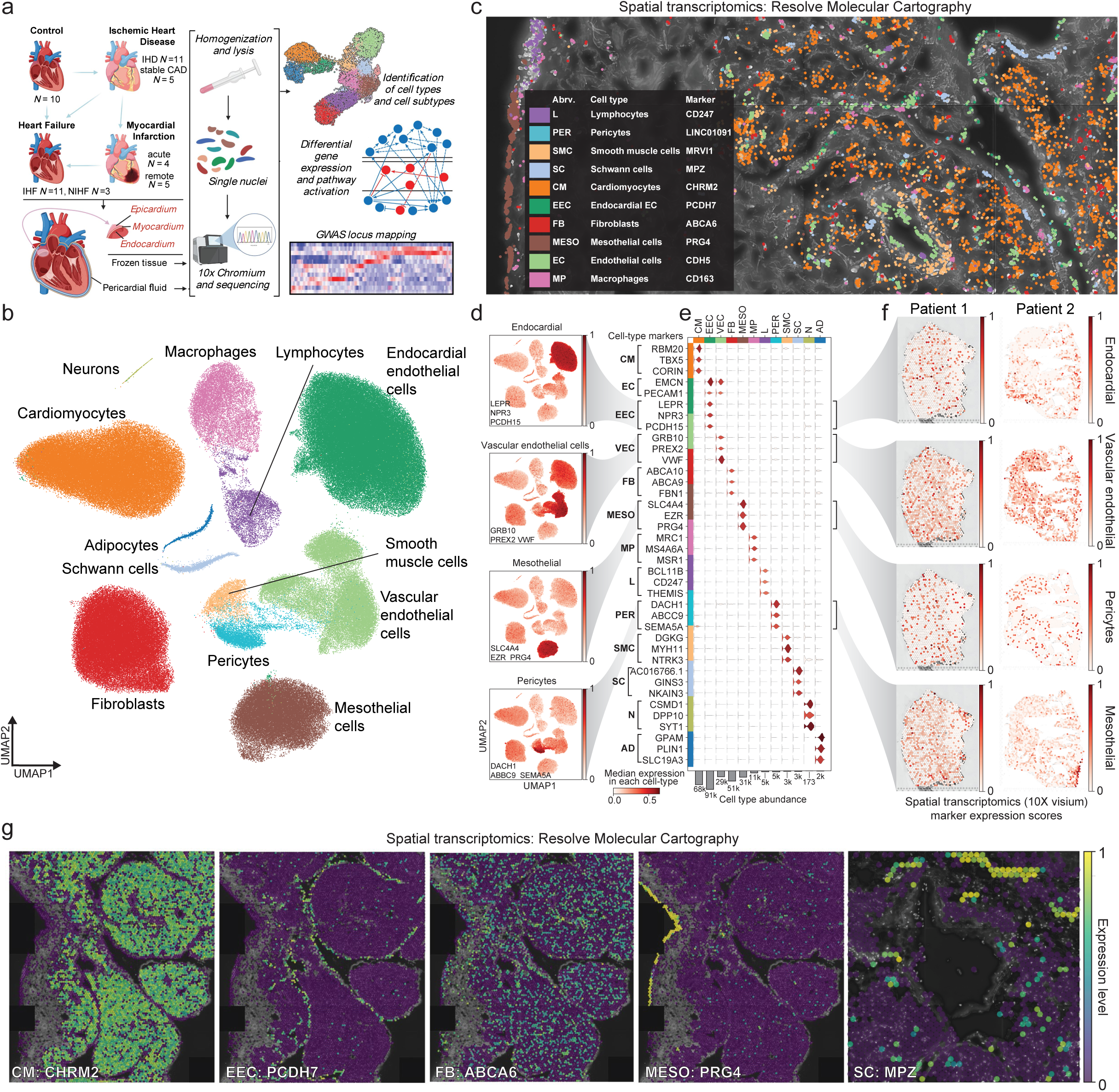
Expression map of the *ex vivo* tissue from the right atrium. **a.** Overview of the samples and experimental approach. Image was created using BioRender. **b.** UMAP embedding of the snRNA-seq data distinguishing populations of 12 main cell types. **c.** Spatial transcriptomics image (Resolve Biosciences) of the cardiac tissue, depicting main cell types in different colors as marked by the expression of their marker genes. **d.** UMAP embedding of the snRNA-seq data highlighting the expression of the marker genes for endocardial endothelial cells (*LEPR*, *NPR3*, and *PCDH15*), vascular endothelial cells (*GRB10*, *PREX2*, and *VWF*), epicardial mesothelial cells (*SLC4A4*, *EZR*, and *PRG4*), and pericytes (*DACH1*, *ABCC9*, and *SEMA5A*). **e.** Violin plot of the marker genes from (d) and other main cell types from (b) (Cardiomyocyte (CM), endothelial cell (EC), endocardial EC (EEC), vascular EC (VEC), fibroblast (FB), epicardial mesothelial cell (MESO), macrophage (MP), lymphocyte (L), pericyte (PER), smooth muscle cell (SMC), Schwann cell (SC), Neuron (N), adipocyte (AD)). **f.** Spatial expression (Visium, 10x Genomics) of the marker genes for endocardial endothelial cells (*LEPR*, *NPR3*, and *PCDH15*), vascular endothelial cells (*GRB10*, *PREX2*, and *VWF*), epicardial mesothelial cells (*SLC4A4*, *EZR*, and *PRG4*), and pericytes (*DACH1*, *ABCC9*, and *SEMA5A*) in two patient samples. **g.** Spatial expression (Resolve Biosciences) of the cardiomyocyte (CHRM2), endocardial endothelial cell (PCDH7), fibroblast (ABCA6), mesothelial cell (PRG4), and Schwann cell (MPZ) marker genes in the right atrial tissue.

The distribution of detected cell types corresponded to expected ratios in the cardiac tissue (*1–9*), given the collection of all three layers of the heart wall – epicardium, myocardium, and endocardium (**Fig. 1a** and **Fig. S1b**) – with cardiomyocytes (CM, 23%), endocardial endothelial cells (EEC, 30%), vascular endothelial cells (VEC, 10%), fibroblasts (FB, 17%), and mesothelial cells (MESO, 10%) as the main cell types (**Fig. 1b** and **Fig. S1c**).

Further examination of the marker genes revealed several candidates that could help to identify the different cell types in spatial transcriptomics (**Fig. 1c**, **f-g**, and **website,** *see* **Fig. S1d** *for instructions*). For example, endocardial endothelial cells expressed several genes that allowed their distinction from vascular endothelial cells, including *LEPR*, *NPR3*, and *PCDH15*, whereas vascular endothelial cells expressed *GRB10*, *PREX2*, and *VWF* (**Fig. 1d-f**). We confirmed several of the marker genes with spatial transcriptomics, using a low-resolution technique (Visium, 10X Genomics) that captures 1-10 cells per spot (**Fig. 1f**), and a high-resolution technique (Molecular Cartography^TM^, Resolve Biosciences) to reach sub-cellular resolution (**Fig. 1c** and **1g**), establishing specific marker genes for different cell types, such as *CHRM2* for cardiomyocytes, *PCDH7* for endocardial endothelial cells, *ABCA6* for fibroblasts, *PRG4* for epicardial mesothelial cells, and *MPZ* for Schwann cells (**Fig. 1g**). Remarkably, visualization of some of the best cellular marker genes identified in snRNA-seq, such as *NPR3* for endocardial endothelial cells, failed due to low abundance and specificity issues in the high-resolution spatial mapping (**Fig. 1e and Fig. S1e**).

### High-definition map of the right atrial vasculature

To identify vascular cell types and cell subtypes, we used a combination of known and novel marker genes distinguishing a total of 11 populations, many of which were previously incompletely recovered (**Fig. 2a-c** and **Fig S1f-g**) (*1-3,26*). We utilized the identified marker genes in spatial transcriptomics to characterize the cardiac vasculature – coronary arteries, arterioles, arterial and venous capillaries, and veins – in our samples (**Fig. 2d-g**) and confirmed the localized expression of *KLF2* into the vascular endothelial cells (**Fig. 2h).** KLF2 is a mechano-activated transcription factor that integrates hemodynamic and proinflammatory stimuli to maintain vascular homeostasis and integrity (*27, 28*). Notably, in snRNA-seq, most of the vascular endothelial subtypes showed downregulation of *KLF2* in the IHD group compared to control samples (**Fig. 2i**), indicating the emergence of endothelial cell dysfunction in the cardiac vasculature, which has been shown to be critical in the initiation and progression of cardiovascular disease (*29*).

**Fig. 2.**
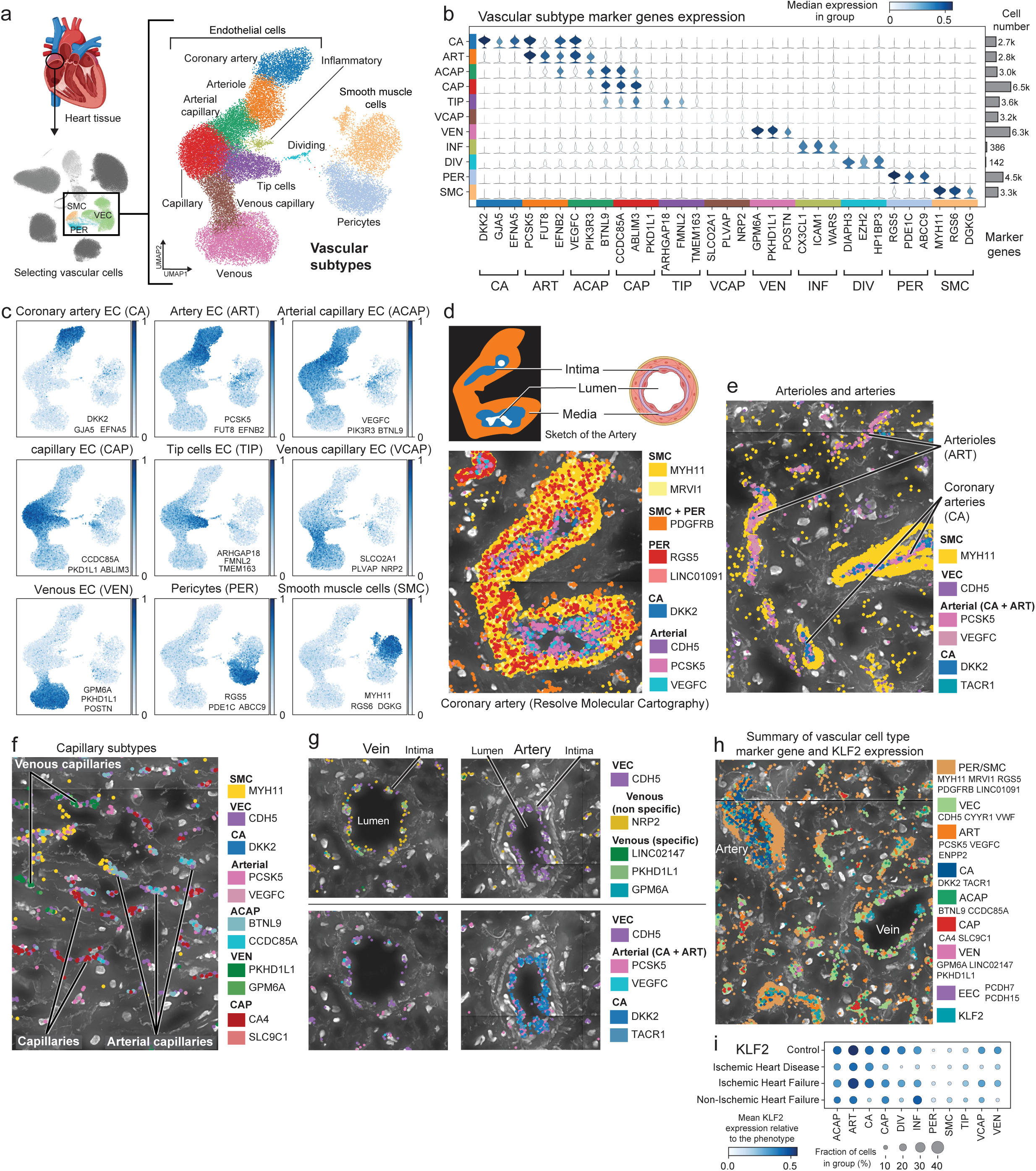
High-definition map of the right atrial microvasculature. **a.** UMAP embedding of the snRNA-seq data for vascular cell subtypes, separating distinct endothelial, smooth muscle cell, and pericyte populations. Heart image from BioRender. **b.** Marker genes for vascular subtypes (coronary artery (CA), arteriole (ART), arterial capillary (ACAP), capillary (CAP), tip cells (TIP), venous capillary (VCAP), venous (VEN), inflammatory ECs (INF), dividing ECs (DIV), pericytes (PER), and smooth muscle cells (SMC)). **c.** UMAP embeddings showing vascular marker gene expression for coronary arteries (*DKK2*, *GJA5*, and *EFNA5*), arterioles (*PCSK5*, *FUT8*, and *EFNB2*), arterial capillaries (*VEGFC*, *PIK3R3*, and *BTNL9*), capillaries (*CCDC85A*, *ABLIM3*, and *PKD1L1*), tip cells (*ARHGAP18*, *FMNL2*, and *TMEM163*), venous capillaries (*SLCO2A1*, *PLVAP*, and *NRP2*), venous ECs (*GPM6A*, *PKHD1L1*, and *POSTN*), pericytes (*RGS5*, *PDE1C*, and *ABCC9*), and smooth muscle cells (*MYH11*, *RGS6*, and *DGKG*). **d- g.** Spatial images (Resolve Biosciences) of coronary arteries and arterioles (**d-e**), venous and arterial capillaries (**f**), and a vein and an artery (**g**) in right atrial tissue. Cross-section of an artery from BioRender. **h.** Summary image (Resolve Biosciences) of different vessels and *KLF2* expression in them. **i.** *KLF2* expression in snRNA-seq data across disease groups in vascular cell subtypes.

### Disease drives inflammatory microvascular dysfunction in the right atrium

To dissect similarities and differences in cardiac cell function on pathway level across main cell types in the three groups (IHD, IHF, and NIHF), we determined the differentially expressed genes across all cell types and cell subtypes using Nebula (*30*) and interpreted the results using Ingenuity Pathway Analysis (IPA, Qiagen) (*31*) (**Fig. 3a**, *data available at* **website**). Among the most significant enrichments in IHD and IHF, were fibrosis for fibroblasts (e.g., ‘Rho’, ‘RHOGDI’ and ‘Apelin’ signaling) and barrier disruption and hypertrophy for endocardial cells (e.g., ‘thrombin’, ‘adherens junction’, ‘cardiac hypertrophy’, and ‘NFAT in cardiac hypertrophy’ pathways). In addition, both fibroblasts and endocardial endothelial cells were enriched for growth factors (e.g., ‘HGF signaling’) and mesenchymal transition (e.g., ‘regulation of the Epithelial to Mesenchymal Transition (EMT) pathway’ and ‘regulation of the EMT by growth factors’), suggesting differentiation/activation of dysfunctional phenotypes, such as myofibroblasts (*32*), and transition to pathological state, such as endocardial dysfunction (*33*), that in early stage disrupts normal tissue function and later on contributes to progression of heart failure.

**Fig. 3.**
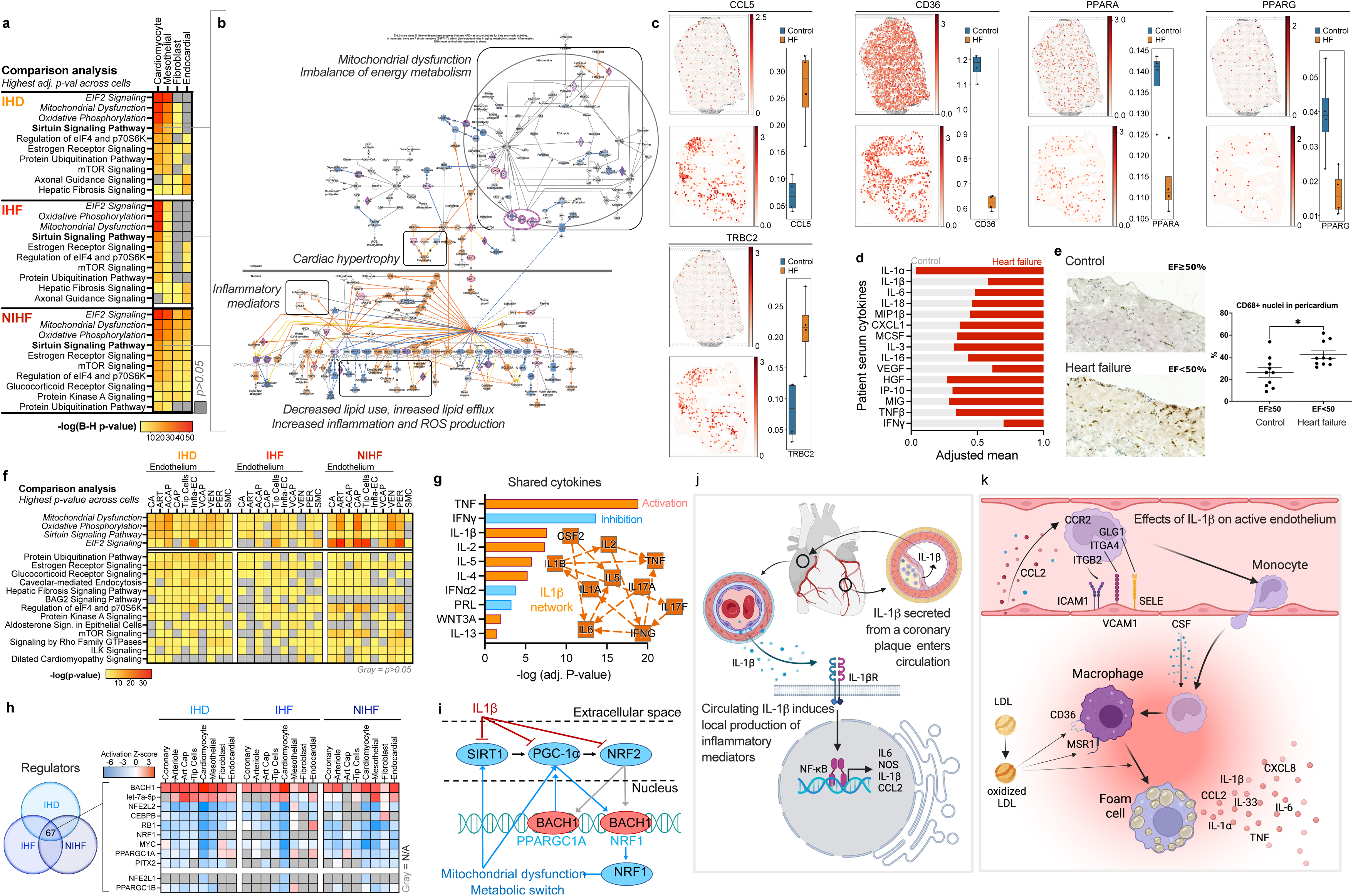
Cardiovascular disease drives inflammatory microvascular dysfunction in the right atrium. **a.** Top 10 canonical pathways with the highest significance score (B-H adjusted p-value p<0.05 for all) by IPA’s comparison analysis across the main cell types in IHD (*N*=11), IHF (*N*=11), and NIHF (*N*=3) against control (*N*=6). Full table can be found at **website**. **b.** Sirtuin signaling pathway from IPA for cardiomyocytes in NIHF against control as a representative for all three groups. Blue indicates inhibition of the molecule, orange/red activation. Purple highlight marks significantly differentially expressed molecules in the dataset. **c.** Spatial expression (Visium) of *TRBC*, *CCL5*, *CD36, PPARA*, and *PPARG* in a control and heart failure sample. Quantitation of the signal across control (*n*=4) and heart failure (*n*=4) sections is shown. Whiskers show the maximum and minimum values, except for outliers (more than 1.5 times the interquartile). **d.** Scaled (0–1) mean of serum cytokines in patients with heart failure (*N*=6) and their controls (*N*=7). No statistical significance. **e.** Quantitation of CD68+ tissue macrophages in pericardium of patients with heart failure (*N*=10) and their control group (*N*=10) (*p<0.0104). A representative immunohistostaining image provided for both groups, showing macrophages in brown. **f.** Top 10 canonical pathways with the highest significance score (Fisher’s Exact Test, p<0.05 for all) across vascular cells within each group: IHD, IHF, and NIHF. Top 10 pathways for each condition were selected and p-value is presented across all conditions. **g.** Top shared cytokines in IPA’s Upstream Regulator analysis between IHD, IHF, NIHF, and disturbed flow in HUVECs dataset (*N*=3 for d-flow and for control). IL-1b network from 2h time point by IPA’s machine learning based graphical summary is a representative for all four time points (2, 8, 14, 32h of IL-1b treatment compared to control (N=3) in HAECs) **h.** Top overlapping Upstream Regulators (Fisher’s Exact Test p<0.05 for all) with similar activity patterns from IPA (microRNAs and transcriptional regulators only) for IHD, IHF, and NIHF in main cell types and four EC subtypes. Red is for predicted activity and blue for inhibition. **i.** Depiction of the connections between the potential mediators of the mitochondrial and metabolic changes. **j.** Overview of IL-1b signaling. **k.** Effects of IL-1b on activated endothelium. j-k. were created with BioRender.

The top pathways in cardiomyocytes and epicardial mesothelial cells were the same across all three comparisons, i.e., ‘EIF2 signaling’ ‘mitochondrial dysfunction’, ‘oxidative phosphorylation’, and ‘sirtuin signaling’. A closer look into sirtuin pathway in cardiomyocytes across the three comparisons (IHD, IHF, and NIHF), revealed similarity across all conditions (**Fig. 3b** and **Fig. S2a**), suggesting accumulation of lipids, lipid peroxidation, and inflammation in the tissue due to impaired mitochondrial function, metabolic changes, cardiac hypertrophy, and increased production of reactive oxygen species (ROS) and inflammatory mediators. To test these predictions in the IHF and control samples, we utilized spatial transcriptomics (**Fig. 3c** and **Fig. S2b**). Compared to control, the IHF tissue exhibited increased expression of many pro-inflammatory mediators, such as *TRBC2* – an immune response activator (*34*), and *CCL5* – a chemoattractant for blood monocytes, memory T helper cells and eosinophils (*35*). In addition, the expression of key metabolic mediators and regulators, such as CD36, which mediates the uptake of long-chain fatty acids in cardiac tissue, and its regulator, nuclear receptor peroxisome proliferator-activated receptor alpha (PPARα), was reduced in IHF compared to control. Impaired synthesis of CD36 has been shown to shorten myocardial energy supply, resulting in insufficient fatty acid uptake and accumulation of toxic lipids, ultimately leading to heart failure (*36–38*). Consistent with the predictions, the expression of the key regulator of fatty acid and energy metabolism, peroxisome proliferator-activated receptor gamma (PPARγ) (*39*), was also decreased in the IHF compared to control sections (**Fig. 3c** and **Fig. S2b**). Cytokine screening from the patient serum samples confirmed the presence of many pro-inflammatory cytokines (**Fig. 3d**), including IL-1-associated cytokines, such as IL-1*α*, IL-1*β*, IL-6, IL-18, MIP1*β*, and CXCL1, chemoattractants and immune cell activators, such as IL-3, and IL-16, growth factors, such as MCSF, HGF (increased in HF) and VEGF (decreased in HF), and biomarkers for heart failure and left ventricular dysfunction, such as IP-10 (CXCL10, increased in HF), and MIG (CXCL9, increased in HF) (*40*). IL-1 has been shown to contribute to all stages of atherosclerotic plaque life from initiation to rupture, although circulating IL-1 levels in patients rarely correlate with disease severity (*41*). For example, circulating levels of IL-1*α* and IL-1*β* can be below detection level even in high-risk patients, yet their neutralization reduces adverse effects and improves disease outcomes (*42, 43*). Here, IL-1*α* exhibited increased and IL-1*β* decreased presence in HF group compared to control, although sample numbers were too small for statistical significance (**Fig. 3d**). Taken together, the results suggested cytokine-induced pro-inflammatory activation and resulting accumulation of immune cells close to vasculature, endocardium, and epicardium in HF samples, which was confirmed with immunostaining of CD68+ tissue macrophages from HF and control samples (**Fig. 3e**).

As suggested by the serum cytokine levels (**Fig. 3d**), production of a potent angiogenic factor, VEGF (*44*), was found to be suppressed in cardiac tissue cells in all three conditions (IHD, IHF and NIHF), although induction of its regulator, hypoxia-inducible factor (HIF1*α*) was observed expectedly in ischemic (IHD and IHF) cardiomyocytes (**Fig. S2c**). Similar trend was observed across endothelial cell subtypes suggesting inhibition of VEGF-regulated angiogenesis (**Fig. S3**). Hypoxia-induced response patterns indicated endothelial activation across most subtypes and diseases, but endothelial movement and/or blood vessel maturation remained mostly inhibited, suggesting endothelial dysfunction and abnormal angiogenesis (**Fig. S4**). Similarly, consistent with the serum cytokine levels and spatial transcriptomics results (**Fig. 3c-d**), CCL2-, TNF-, and CCL5-induced vasoconstriction was increased across endothelial subtypes and diseases (**Fig. S5**). Taken together, the results suggested microvascular dysfunction with an inflammatory component.

Closer look into the differential expression enrichments in the vascular cells and cell subtypes revealed high similarity with the other main cell types, having the same four pathways (‘mitochondrial dysfunction’, ‘oxidative phosphorylation’, ‘sirtuin signaling’, and ‘EIF2 signaling’), top the most significant enrichments across all three conditions (**Fig. 3f**). To recreate endothelial dysfunction in an *in vitro* model, we exposed Human Umbilical Vein Endothelial Cells (HUVECs) to disturbed flow (d-flow, i.e., oscillatory non-laminar shear stress) (*45*). We compared these changes to changes observed in the human cardiac tissue and saw decreased mitochondrial metabolic activity, and increased HIF1*α* activity, cardiac hypertrophy signaling, and pro-inflammatory changes comparable to those in the tissue cells (**Fig. S6a**). To screen for potential intercellular mediators of the observed changes, we extracted overlapping cytokines from the three conditions (IHD, IHF, NIHF) and d-flow endothelial cells using Upstream Regulator analysis of IPA (**Fig. 3g**), and found several cytokines that were measured from the patient serum, such as TNF, which was more abundant in HF patient samples compared to controls, IFN*γ*, which was more abundant in controls, and general enrichment of IL-1-related factors (**Fig. 3d**). The analysis suggested IL-1*β* as a central regulator of the observed gene expression changes and a driver of the observed cytokine expression. To validate this finding, we performed RNA-seq from IL-1*β*- stimulated Human Aortic Endothelial Cells (HAECs), confirming an IL-1*β*-centric interleukin network connecting the observed mediators in the cells (**Fig. 3g** and **Fig. S6b**). To further identify the potential regulators of the shared functional changes across cardiac cell types, we overlapped transcriptional regulators and microRNAs from Upstream Regulator analysis of IPA for the three conditions (IHD, IHF, and NIHF) (**Fig. 3h**). Among the top regulators were transcription factors BACH1, NRF2 (*NFE2L2*), and NRF1 and a co-activator PGC1*α* (*PPARGC1A*) (**Fig. 3h**), which are known regulators of mitochondrial function, metabolism, and cellular redox stress (*46–49*). Their interactive network (**Fig. 3i**) may thereby partially explain the observed pathway changes in the main cell types. The notion was supported by the data collected from IL-1*β*-stimulated HAECs (**Fig. S6c**), which illustrated a switch to anaerobic glycolysis, downregulation of mitochondrial biogenesis and metabolism, increased inflammation, ROS production, and hypertension signaling, highlighting suppression of SIRT1, PGC1*α*, and NRF2 as mediators of these events (**Fig. 3i**). Collectively, these data suggest that IL-1*β*, which has been shown to be a component of coronary plaques (*50*) and produced by activated endothelia, smooth muscle cells, fibroblasts, and immune cells in the body (*41*), travels to RA and induces local production of inflammatory mediators (**Fig. 3j**) and recruitment of immune cells (**Fig. 3k**). Accumulation of immune cells (**Fig. 3c** and **3e**), lipids (**Fig. 3b**), and oxidative stress (**Fig. 3b**) are known predecessors of foam cell formation (**Fig. 3k**) (*51*).

### Disease-driven immune cell accumulation causes chronic inflammation in the atrium

To explore the inflammatory changes in the right atrium, we next characterized the immune cell populations (**Fig. 4a**) using previously established marker genes (**Fig. 4b**) (*1-3,52-56*) and GO term enrichments (**website**). The immune clusters consisted of rich populations of macrophages (MPs) and T cells, in addition to dendritic cells, B cells, and plasma cells. Spatial transcriptomics of the atrial tissue illustrated immune cell clusters that localized to epicardial mesothelium and vascular endothelium (**Fig. 4c**). The most prevalent immune cell population across all disease groups was macrophages (**Fig. 4d**). Other subtypes displayed more variability, suggesting increase in abundance from control to more severe disease (**Fig. 4d**). In spatial analyses, MP marker gene, *CD163,* co-localized with Infla-MP marker gene, *LYVE1,* with increased expression in IHF compared to control, consistent with previous findings (**Fig. 3c** and **3e**), whereas LT-CD4 marker gene, *IL7R,* expression overlapped only partially with the MP marker genes, although similarly higher expression in IHF sections compared to control was observed (**Fig. 4e** and **Fig. S7a**).

**Fig. 4.**
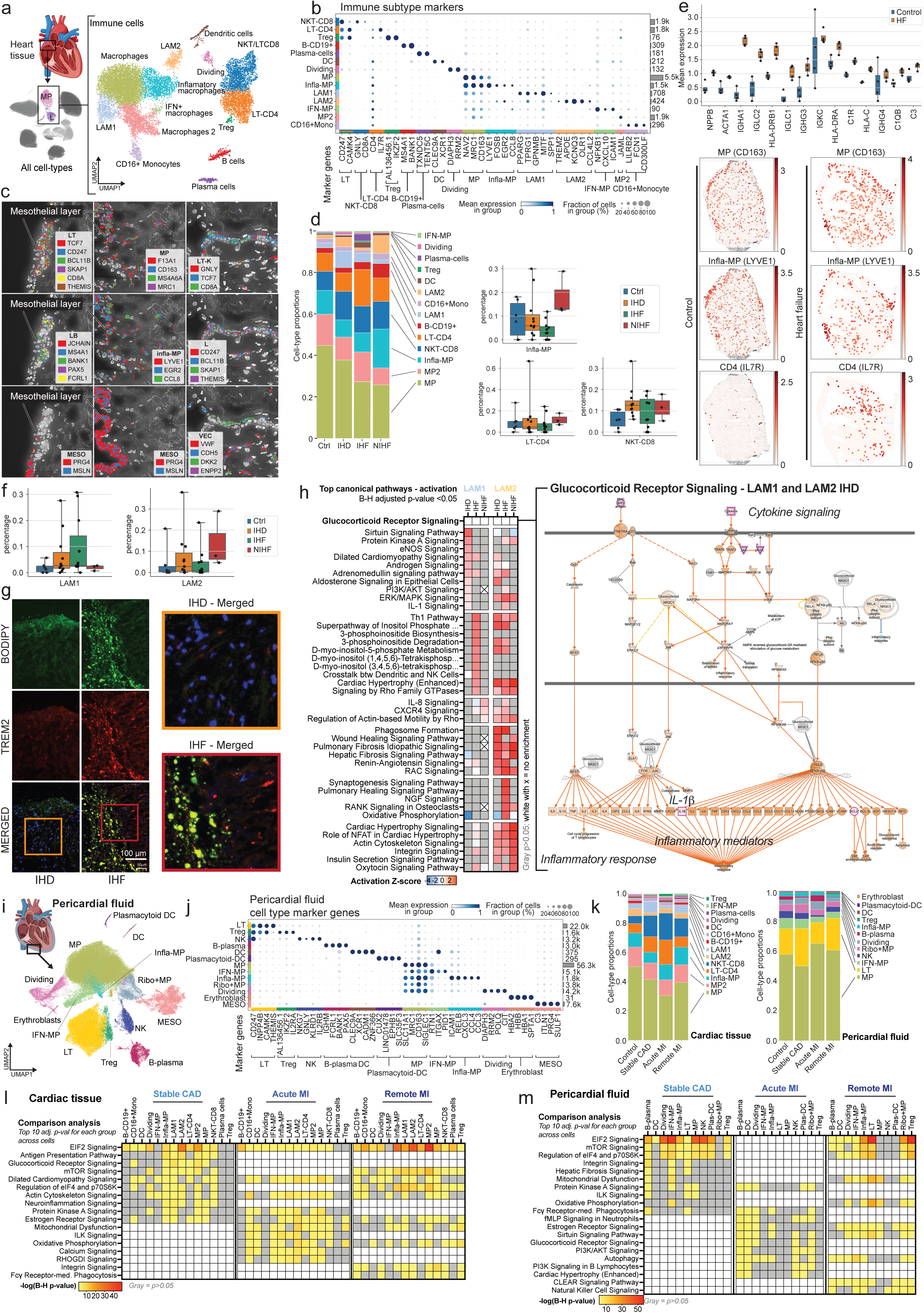
Disease-associated changes in the immune cells of the right atrial tissue and the pericardial fluid. **a.** snRNA-seq for immune cell subtypes in right atrial tissue, separating 14 populations. Heart image from BioRender. **b.** Marker genes for immune subtypes (dendritic cells (DC), macrophages (MP), lipid-associated macrophages (LAM), interferon MP (IFN-MP), monocytes (mono). **c.** Spatial expression images (Resolve Biosciences) of the cardiac tissue, depicting immune cells as marked by the expression of their marker genes near epicardial mesothelium and vasculature. **d.** Immune cell proportions in control, IHD, IHF, and NIHF samples. Bar charts depict the immune cell subtype proportions (%) of all cells in each sample for LT-CD4, infla-MP, and NKT-CD8 in control, IHD, IHF, and NIHF groups. Whiskers show the maximum and minimum values, except for outliers (more than 1.5 times the interquartile). **e.** Quantitation of the spatial expression signal (Visium) for *NPPB*, *ACTA1*, and immunoglobulins in control (*n*=4) and heart failure (*n*=4) tissue sections. Representative Visium images shown for *CD163*, *LYVE1*, and *IL7R* expression. Whiskers show the maximum and minimum values, except for outliers (more than 1.5 times the interquartile). **f.** LAM1 and LAM2 proportions (%) of all cells in each control, IHD, IHF, and NIHF sample depicted by groups. Whiskers show the maximum and minimum values, except for outliers (more than 1.5 times the interquartile). **g.** Immunofluorescence images of lipid-associated macrophages (LAM) labeled using anti-TREM2 (red) and bodipy (green) in the epicardial side of the human right atrial appendage. Patients with ischemic heart failure (IHF; middle panel) had increased number of LAMs compared to patients with ischemic heart disease (IHD; left panel). Right panel shows zoomed region of interest from the merged image. Representative samples are shown. **h.** Top canonical pathways with highest activation Z-scores for LAMs in IHD (*N*=11), IHF (*N*=11), and NIHF (*N*=3) against control (*N*=6) based on IPA analysis. Predicted pathway activation is shown in red and inhibition in blue based on the expression changes of the pathway molecules in the dataset and current literature-based knowledge curated into the QIAGEN Knowledge Base. Gray is for enriched pathways with B-H adjusted p-value over 0.05 and white with x for no enrichment. All pathways with activation Z- score are statistically significant (B-H adjusted p-value < 0.05). A representative Glucocorticoid receptor pathway chart is shown. **i.** UMAP embedding of the snRNA-seq data for immune cell subtypes in pericardial fluid, separating 12 immune cell populations and a mesothelial population. Heart image by BioRender. **j.** Marker genes for immune subtypes in the pericardial fluid. **k.** Immune cell proportions in cardiac tissue and corresponding pericardial fluid samples in control, stable CAD, acute MI, and remote MI. **l-m.** Top 10 pathways with highest significance score in IPA’s comparison analysis (B-H adjusted p-value < 0.05) shown across the cell types in each condition in cardiac tissue (**l**) and corresponding pericardial fluid samples (**m**) in stable CAD (N=5), acute MI (N=4), and remote MI (N=5) compared to control (N=4). Only the top 10 for each group are shown. Full data can be explored at **website**.

Chronic inflammatory response is part of adverse cardiac remodeling that precedes heart failure development (*57*). This remodeling is characterized by re-expression of the fetal gene program that includes natriuretic peptide B (NPPB) and α 1 skeletal muscle actin (ACTA1) (*58*), both of which were observed in the IHF sections (**Fig. 4e** and **Fig. S7a**). To a lesser amount, the genes were also expressed in the control sections and colocalized to the same areas of the section, indicating disease-associated changes in gene expression before clinical manifestation of the disease. However, strong immunoglobulin expression was only observed in the IHF, consistent with chronic inflammatory state and accumulation of antibody deposits in the failing atrial myocardium.

As a confirmation for the results in the microvasculature that suggested lipid accumulation and early stages of foam cell formation (**Fig. 3**), we detected two populations of Lipid associated macrophages (LAMs), out of which LAM1 indicated partiality to ischemic disease and LAM2 to NIHF (**Fig. 4f**). We confirmed the colocalization of an established LAM marker TREM2 (*59*) with lipid droplets in IHF sections and observed increased signal in IHF compared to IHD (**Fig. 4g**).

Exploration of pathway enrichments for differentially expressed genes in the LAM populations revealed differences in the pathway numbers, their sharing, and in the enriched pathways themselves. Altogether, the LAM1 population enriched for 118 and LAM2 for 228 pathways (B-H adjusted p-value < 0.05). 10% of the LAM1 pathways were not found from the LAM2 populations, including ‘nitric oxide’, ‘inositol phosphate’ and ‘phosphoinositide signaling’ related terms, and ‘circadian rhythm’, ‘ferroptosis’, and ‘senescence signaling’ pathways (**Data S1**). In IHD, the top activated pathways (based on activation Z-score by IPA) were ‘sirtuin signaling pathway’, ‘protein kinase A signaling’, ‘eNOS signaling’, and ‘dilated cardiomyopathy signaling’ for the LAM1 population and ‘cardiac hypertrophy (enhanced)’, ‘phagosome formation’, ‘wound healing’ and ‘fibrosis signaling’ pathways for LAM2. Half of the enriched LAM1 pathways were shared with LAM2, including pathways for ‘oxidative phosphorylation’, ‘mitochondrial dysfunction’, ‘sirtuin signaling’, ‘estrogen receptor signaling’, and ‘glucocorticoid receptor signaling’, comparably with the main cell types (**Fig. 3a, 3f,** and **Data S1**). ‘Glucocorticoid receptor signaling’ was among the most significant enrichments in all but one LAM group (LAM2 – IHF) (**Fig. 4h** and **website**), highlighting increased cytokine-mediated production of IL-1*β*, among other key pro-inflammatory mediators. The LAM1-specific pathways included ‘IL-1’ and ‘IL-6 signaling’ among others, whereas LAM2-specific populations consisted of ‘neuronal’, ‘inflammatory’, ‘cardiac hypertrophy’, and ‘growth factor’ related pathways (**Data S1**). Only 34% of the LAM1 pathways in IHD were shared with the LAM1 pathways in IHF and none with NIHF (**Fig. 4h** and **Data S1**). LAM1s in NIHF only enriched for 11 pathways altogether, seven of which were shared with IHF. LAM2 populations shared more pathways between conditions (**Fig. 4h** and **Data S1**). For example, only 13% of the IHD-enriched pathways were unique to IHD, 27% were shared with IHF, 21% with NIHF, and 39% were shared by all three conditions. Overall, LAM2 in IHF enriched for more pathways than the other two conditions and therefore also had more unique pathways. Collectively, the pathway enrichments in the LAM1 population consisted mostly of ‘extracellular signaling’, ‘metabolism’, and ‘senescence’ related terms, and in the LAM2 population, of pathway terms that intersect with ‘cardiovascular disease’, ‘inflammation’, and ‘cancer’, suggesting response to environmental cues for LAM1s and pro-disease signaling for LAM2s.

### Pericardial fluid cells reflect the changes in disease states

Pericardial fluid is an enriched milieu of cytokines, growth factors, and cardiac hormones that reflect and regulate overall heart function. Despite the diagnostic and therapeutic potential, its cell composition has not been thoroughly examined with single-cell techniques (*60*) and matching tissue and fluid samples have not been explored. Given the detected accumulation of immune cell clusters to the epicardial mesothelium in spatial transcriptomics (**Fig. 4c, 3c and 3e**), we next decided to explore the cell populations in paired right atrial tissue and pericardial fluid samples of the IHD patients to gain insight into the potential interactions between the tissue and the fluid. Although the tissue biopsy comes from a specific site, the pericardial fluid is not specific to only one anatomical region but provides a more integrated view of the cardiac function.

We focused on three different groups: patients with stable coronary artery disease who underwent elective surgery (stable CAD), patients with acute myocardial infarction (acute MI), and patients, who had suffered infarction earlier in life (remote MI) to compare stable disease with acute phase and recovery from the acute phase. Using established marker genes, pericardial fluid was, expectedly, found to be enriched for immune cells (**Fig. 4b** and **4i-j**). The cell proportions between the tissue and the fluid showed both similarities and differences (**Fig. 4k**). The largest population in both was macrophages, followed by lymphocytes and natural killer cells. Cardiac tissue-specific populations included LAMs, monocytes, and plasma cells, and pericardial fluid included populations of Ribo+MPs (i.e., MPs with increased expression of ribosomal genes), plasmacytoid dendritic cells (plas-DCs), and erythroblasts that were not detected in the cardiac tissue. Infla-MPs were proportionally larger population in the tissue compared to the fluid, whereas IFN-MPs were more predominant among the fluid immune cells.

In the tissue samples, MP2 population seemed to be more prevalent in the control samples, whereas NKT-CD8 and infla-MPs were more abundant in stable CAD and MI groups (**Fig. S7b**). LT-CD4 population was more numerous in the stable CAD and acute MI, but in remote MI the levels resembled control levels. In the pericardial fluid, MP, LT, infla-MP, and Treg proportions were more variable in the IHD groups compared to control group, suggesting active exchange between the tissue and the fluid (**Fig. S7b**).

Differentially expressed gene enrichments were similar, although not identical, in the pericardial fluid and corresponding tissue with 66-68% of all enriched pathways being shared between the fluid and tissue cells in each of the three groups (**Fig. 4l-m** and **website**). Stable CAD and remote MI had 100% overlap in pathways enriched in each, both in fluid and tissue samples, although the rank order was a mix of stable CAD and acute MI in remote MI, likely due to its shared disease component with both groups (**Fig. 4l-m, Data S1,** and **website**). 12% and 18% of enriched pathways were unique to acute MI in fluid (63 unique pathways) and tissue (83 unique pathways), respectively, including terms related to ‘IL-1 signaling’ and ‘metabolism’ in the fluid and ‘secondary messenger signaling’, ‘interleukins’, and ‘metabolism’ in the tissue. In tissue, macrophages were the most active populations in all three conditions, promoting ‘immune response’, ‘intracellular and second messenger signaling’, and ‘cellular stress’ and ‘injury’ in stable CAD and ‘cardiovascular signaling’ and disease-specific pathways in acute MI, leading to glucocorticoid-mediated CCL2 and CCL3 production in both (**Fig. S7c-d**). CCL2 is a regulator of macrophage recruitment and polarization in inflammation (*61*) and CCL3 is an acute pro-inflammatory recruiter and activator of leukocytes in the heart tissue, as well as an inducer of TNF and IFN production that is associated with cardiomyocyte injury, cardiac dysfunction, and delayed ventricular repolarization (*62*).

To take a closer look into the pericardial fluid cells, we chose one of the most prominent specialized populations in the ischemic patients, IFN-MPs (**Fig. 4k**). Dissection of the co-expression patterns in the IFN-MP population highlighted several functional modules, such as ‘lymphocyte activation’, ‘cell migration’, ‘cytokine and interferon signaling’, ‘oxidative phosphorylation’, and ‘cholesterol metabolism’ (**Fig. 5a** and **website**). The proportion of the cells was generally lower in the tissue of stable CAD and MI patients and higher in their pericardial fluid, suggesting mobilization of the cells in the earlier and acute stages of the disease, whereas in the more advanced disease, the data suggested potential accumulation in the tissue (**Fig. 5b**). In spatial images (Visium), *CXCL10*, a marker gene for the IFN-MP population, was detected in IHF, clustering in the myocardium, but not in the control sections (**Fig. 5c** and **Fig. S8a**). CXCL10 (also known as interferon gamma-induced protein 10, IP-10) is a chemoattractant and polarizing factor for various immune cells, promoter of T cell adhesion to ECs, inhibitor of angiogenesis and a biomarker for heart failure and left ventricular dysfunction (**Fig. 3d**) (*40, 63,64*).

**Fig. 5.**
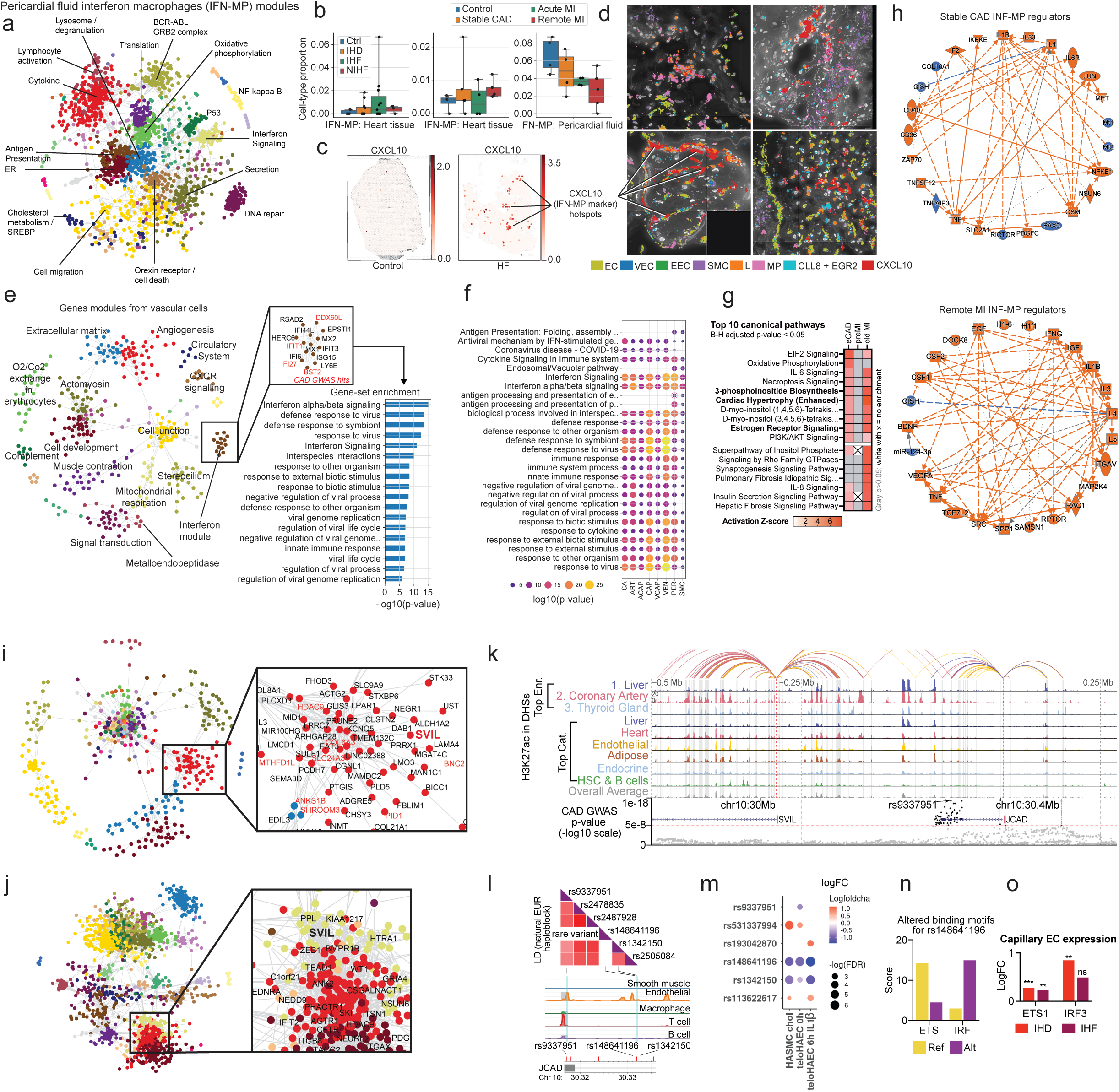
Disease-associated genetic variation affects disease-relevant modules across cell types. a. Gene expression modules in IFN-MPs of the pericardial fluid. **b.** IFN-MP proportions (%) of all cells in control, IHD, IHF, and NIHF samples and tissue and pericardial fluid control, stable CAD, acute MI and remote MI samples depicted by group. Whiskers show the maximum and minimum values, except for outliers (more than 1.5 times the interquartile). **c.** Representative Visium images shown for *CXCL10* expression. **d.** Spatial images (Resolve) for EC (*EMCN, ERG, PECAM1, CDH5, VWF*), VEC (*DKK2, ENPP2, PCSK5, CYYR1*), EEC (*PCDH7*), SMC (*NTRK3, MRVII), L (BCL11B, CD247, SKAP1, THEMIS*), MP (*CD163, MRC1, F13A1, MS4A6A*), inflammatory (*CCL8* and *EGR2*), and *CXCL10* gene expression. **e.** Gene expression modules for vascular cells (all subtypes combined) highlighting interferon module, its genes (GWAS-linked genes in red), and pathway enrichments. **f.** Enrichments of the interferon module across vascular cells. **g.** Top canonical pathways with highest activation Z-scores for IFN-MPs of the pericardial fluid in stable CAD (N=5), acute MI (N=4), and remote MI (N=5) compared to control (N=4) based on IPA analysis. Predicted pathway activation is shown in red based on the expression changes of the pathway molecules in the dataset and current literature knowledge curated into the QIAGEN Knowledge Base. Gray is for enriched pathways with B-H adjusted p-value over 0.05 and white with x for no enrichment. All pathways with activation Z-score are statistically significant (B-H adjusted p-value < 0.05). **h.** Regulator networks for stable CAD and remote MI in IFN-MPs from IPA’s graphical summary. **i-j.** Gene expression module for SVIL in SMCs **(i)** and pericardial fluid cells **(j)**. GWAS-linked genes are shown in red. **k.** Epimap linking of the *JCAD/SVIL* locus. **l.** Illustration of LD SNPs of the natural European haploblock combined with the scATAC-seq data from human coronary arteries generated using LDlink**. m.** Allele-specific enhancer activity measured with STARR-seq in teloHAECs under basal and inflammatory conditions (6h IL-1b) and HASMCs subjected to cholesterol loading for 24h. SNPs demonstrating significant changes (FDR < 0.1) in enhancer activity are shown. **n.** Transcription factor binding motifs altered by rs148641196. Position weight matrix scores shown for reference and alternate. **o.** Changes in transcription factor (TF) *ETS1* and *IRF3* gene expression in capillary endothelial cells in IHD and IHF in snRNA-seq by Nebula.

In the higher-resolution spatial images, CXCL10 clusters were found in the vicinity of vasculature and endocardium, colocalizing with macrophages, lymphocytes, and the marker genes for infla-MPs (*CLL8* and *EGR2P*) (**Fig. 5d** and **Fig. S8b**). In the vascular cells, module analysis confirmed the presence of an IFN module comprising several IHD-GWAS genes (**Fig. 5e**). The IFN module was present across all vascular cell subtypes and enriched for several pathways linked to ‘antigen presentation’, ‘cytokine signaling’, ‘interferon signaling’, and ‘defense and immune responses’ (**Fig. 5f**), confirming the connection between the vascular cells and the INF-responsive immune cells at the transcriptional level.

Pathway analysis of the IFN-MPs in stable CAD, acute MI, and remote MI revealed high similarity between the stable CAD and remote MI groups, with almost identical pathways being enriched (B-H adjusted p-value <0.05) in both with similar activation Z-scores (**Fig. 5g** and **website**). For example, ‘3-phosphoinositide biosynthesis’, ‘cardiac hypertrophy (enhanced)’ and ‘estrogen receptor signaling’ pathways were among the top pathways in both stable CAD and remote MI. However, there were underlying differences in the pathway molecules. For instance, in the hypertrophy pathway of the remote MI, the analysis highlighted TNF, insulin-like growth factor 1 and pro-inflammatory cytokines as main upstream molecules activating the signaling cascades, whereas in stable CAD, endothelin-1, TNF, and integrins were highlighted (data not shown). Acute MI enriched for similar pathways compared to stable CAD and remote MI, but the activation Z-score was opposite in many cases and only two of the pathways, ‘autophagy’ and ‘AMPK signaling’, were significantly enriched (B-H adjusted p-value < 0.05). Dissection of the biological and disease-associated functions of the IFN-MPs suggested increased ‘blood cell activation’, ‘recruitment’, ‘movement’, ‘adhesion’, ‘migration’, ‘engulfment’, and ‘atherosclerosis’ in stable CAD, decreased ‘lipid droplet accumulation’, ‘blood platelet aggregation’, ‘blood cell accumulation’, and ‘cell movement’ in acute MI, and increased ‘infection’, ‘cell movement’, ‘cell migration’, ‘proliferation’, ‘engulfment’, ‘angiogenesis’, ‘fibrogenesis’, and ‘vascular development’ in remote MI (**Data S1**). Central factors orchestrating these changes included IL-1*β* (stable CAD and remote MI) and IFN*γ* (remote MI) (**Fig. 5h**) among other factors. Taken together, the results indicate that presence of IFN-MPs in the tissue correlates with advanced disease and chronic inflammatory state of the right atrium. Moreover, the data suggests that the IFN-MP population present in the pericardial fluid reflects the changes in the tissue function and responds to environmental cues of acute and chronic disease states.

### Disease-associated genetic variation affects disease-relevant modules across cell types

Genetic studies on cardiovascular disease provide invaluable resources for understanding their causal mechanisms. However, currently discovered loci only explain a small fraction of disease heritability, indicating that additional contributors exist, and new approaches are needed for their discovery. In this study, we sought to dissect IHD-associated GWAS gene interactomes across different cell types using our newly developed module analysis method that combines coordinated gene expression changes across cell types together with GWAS-linked genes, forming functional modules of genes with similar expression patterns (*all data can be explored at* **website)**. One of the genes highlighted by the module analysis was *SVIL* that was present both in the vascular and pericardial fluid cells (**Fig. i-j**) and resides in a GWAS locus on chromosome 10 that gives rise to two IHD-associated genes, *SVIL* and *JCAD*, that link to the same regulatory element bearing several IHD-associated variants (**Fig. 5k**). *JCAD* has been previously implicated in endothelial dysfunction, pathological angiogenesis, and atherosclerotic plaque formation (*65*), whereas the role of *SVIL* in cardiovascular disease remains unknown, but it may promote angiogenesis and epithelial to mesenchymal transition (*66*).

In smooth muscle cells, *SVIL* resided in a module that enriched for ‘glycosaminoglycan’, ‘proteoglycan’, and ‘chondroitin sulfate’ related pathways, whereas *JCAD* module consisted of three genes *JCAD*, *CFAP36,* and *PALMD*, which has been shown to regulate aortic valve calcification via altered glycolysis and inflammation (*67*) (**Fig. 5i** and **Fig. S9a**). Proteoglycans regulate extracellular matrix composition and organization, partake in inflammatory processes through leukocyte adhesion and migration, and are central modifiers of ischemic and pressure overload-related cardiac remodeling that leads to heart failure (*68*). In pericardial cells, *SVIL* module enriched for ‘cell communication’, ‘intracellular signal transduction’, and ‘regulation of cell adhesion’ (**Fig. 5j** and **Fig. S9b**). For example, in T and NK cells the modules enriched for ‘cell signaling’ and ‘communication’, in B-plasma cells for ‘T cell differentiation’, ‘activation’, and ‘signaling’, and in mesothelial cells, for ‘extracellular matrix organization’, ‘blood vessel development’, and ‘morphogenesis of an anatomical structure’ (**Fig. S9c-f**), indicating involvement in several cell-type-specific functions.

Although genome-wide association studies (GWASs) successfully associate genomic loci with diseases, they are restricted in their ability to dissect complex causality, as they do not specify the causal variant(s). For example, the *JCAD/SVIL* locus investigated in this study associates significantly with IHD and several of the top associated variants reside in the regulatory elements that are active in the heart and coronary arteries (**Fig 5i** and **Fig. S10a**). To bridge the gap between GWAS association and biological understanding of underlying disease risk, we utilized our recently published EpiMap analysis (*69*), which suggested rs9337951 as the top candidate (p=1e- 17, **Fig. 5i and 5l**). To further identify individual causal variants, we tested each regulatory variant for its effect on STARR-seq reporter gene expression in HAECs and in primary human aortic smooth muscle cells (HASMCs) subjected to pro-atherogenic stimuli (IL-1*β* and cholesterol- loading). Altogether, six regions of 200 bp were studied representing haplotypes that encompassed nine common variants and seven rare variants in the European population. Of these, two common variants, namely rs9337951 and rs1342150 and four rare variants including rs531337994, rs148641196, rs113622617 and rs193042870 exhibited allele specific enhancer activity (**Fig. 5m** and **Table S1**). To validate the regulatory effect of the identified common variants, we used CRISPR technology to perturb the enhancer activity. First, we performed CRISPR mediated deletion of the regulatory elements in teloHAECs, resulting in 50-70% decrease in *JCAD* expression. This effect was strongest for the rs9337951, the top candidate suggested by the EpiMap analysis, likely due to its exonic location (**Fig. S10b-f**). Unfortunately, *SVIL* expression was not detected in the *in vitro* model despite its significant expression levels in cardiac endothelial cells, highlighting the importance of human tissue data in GWAS dissection (**Fig. S10g**). To ensure no mRNA sequence itself was targeted, we additionally performed CRISPR inhibition experiment where the regulatory elements were targeted with the transcriptional repressor dCas9-KRAB- MeCP2. This approach resulted in similar effects for both targeted regions where on average 70% repression in *JCAD* expression was seen, thus confirming the role of variant-carrying enhancers in gene regulation (**Fig. S10d-f**).

In addition to EpiMap results, motif analysis (*70, 71*) suggested that rs148641196, a rare variant with the highest allele-specific activity in reporter assay (**Fig. 5m**), restored a functional interferon regulatory factor (IRF) binding site from an ETS binding site (**Fig. 5n**) which could potentially promote inflammation-induced dysregulation of the locus during disease development. In our dataset, the expression of both *ETS1* and *IRF3* was upregulated in capillary endothelial cells in IHD and IHF (**Fig. 5o**). ETS1 is an essential factor for vascular angiogenesis (*72*), whereas cardiac damage-induced IRF3-IFN activation has been linked to expression of inflammatory cytokines and chemokines, inflammatory cell infiltration of the heart, and fatal response to myocardial infarction (*73*). Although the rare variant is unlikely to explain general disease-linked signal arising from the *JCAD/SVIL* locus, it suggests that when present, rare variants, such as rs148641196 that are inherited with common GWAS SNPs, may significantly increase individual’s risk for a disease.

Taken together, these data illustrate how a disease-associated variant can affect expression of more than one gene in several cell types, partaking in distinct functions in each through functional gene expression modules. They also show, how less characterized genes can be linked to functions based on their co-expression patterns, which may help to identify novel disease-associated connections beyond previously established pathways.

## Discussion

Chronic cardiovascular diseases cause electrical and structural remodeling of the atrial myocardium, predisposing to sinus node dysfunction and arrhythmias. Here, we provide the first transcriptional and spatial dissection of the *ex vivo* biopsies of the human right atrium in three conditions (IHD, IHF, and NIHF), highlighting evidence of microvascular dysfunction and pro-inflammatory changes in the earlier stages, and subsequent chronic inflammation and hypertrophic signaling in the advanced stages of the disease.

In the healthy human heart, the main source of energy is fatty acids, followed by glucose, and lactate, and to a lesser extent ketone bodies and amino acids (*74*). One of the hallmarks of disease progression is the switch from fatty acids to glucose, and accompanied suppression of mitochondrial oxidative phosphorylation, together with a gradual decrease in mitochondrial biogenesis, and overall downregulation of oxidative metabolism (*74*). As a result, the failing heart reverts to “fetal phase”, where glycolysis is increased, as fatty acid and glucose oxidation are suppressed. However, it remains unresolved whether these metabolic changes cause disease progression or result from it. In this study, we documented the metabolic switch in the earlier stage of the disease prior to major hypertrophic and inflammatory changes of the tissue, across various right atrial cell types from cardiomyocytes to vascular and immune cells. Although metabolic reprogramming would be expected in the hypertrophic tissue of the ventricles, it is more surprising to find it in the right atrial appendage for its remote location, as it suggests major changes in the tissue function far away from the atherosclerotic plaque that presumably causes the ischemic disease. Furthermore, metabolic reprogramming is present in the atrial tissue both in the IHF and NIHF, suggesting underlying mechanistic similarities between the conditions.

As the main causal changes for the disorders studied here are expected to manifest elsewhere, further dissection of the data suggested the pro-inflammatory molecule, IL-1*β*, as the initiator of the metabolic reprogramming and subsequent inflammation in the remote location profiled in this study. IL-1*β* is highly produced and systemically released by activated macrophages, fibroblasts, smooth muscle cells, and endothelial cells in response to danger signals that activate proinflammatory processes, whereas IL-1*α* is a pro-cytokine that is not processed nor released outside the cell, unless the cell is injured, which makes it a marker of cell damage (*75*). In this study, higher levels of IL-1*α* in serum samples of the patients with HF compared to the control samples may have been an indicator of vascular inflammation (*41*), as the gene expression patterns of the cardiomyocytes, mesothelial cells, vascular cells, and various immune cells suggested coronary microvascular dysfunction with an inflammatory component as one of the causal mediators of disease-associated changes in the tissue. Coronary microvascular dysfunction impairs the endothelial lining of the blood vessels that oxygenate and nourish the heart muscle. It manifests as structural and functional abnormalities of the microvasculature and its presence associates with the worst clinical outcomes, especially in myocardial ischemia (*17*). The role this cellular state in the right atrial tissue plays in the disease progression and manifestation of atrial fibrillation and HF warrants further investigation. Although IL-1*β* was identified as an underlying factor promoting the pathological changes more widely, it should be noted that the main regulator may also be one of the factors downstream of IL-1*β* activity, such as IL-6, which has been suggested to play a causal role in IHD (*76*).

Inflammation has long been known to play a detrimental role in the atherosclerotic plaques in IHD (*77*) and to correlate with increased risk for cardiovascular events (*78, 79*), which has prompted the question if inflammation should be targeted in the cardiovascular disease treatment more broadly. As the data presented here suggests an active role for the interleukins in initiation of severe cardiovascular remodeling far from the acute site, targeting low residual inflammation in cardiovascular disease using widely available drugs seems reasonable. Although measurement of IL-1*β* in the clinic is not feasible, C-reactive protein (CRP) is a correlate for IL-1 and IL-6 activity (*80–82*) that is available in every laboratory. As drugs targeting IL-1*β* and IL-6 are available for use in cardiovascular disease treatment (*42,83-87*), further investigation of these interleukins in the disease manifestation and progression in adult human tissue and human disease models is critical.

In conclusion, the integrated information from cell and cell subtype-specific gene expression changes across various cardiovascular traits can be used to better understand the complex molecular disease mechanisms behind the traits, and to identify critical drivers of cardiovascular disease, as well as to guide the development of novel therapeutic strategies for cardiovascular disease treatment. The data collected in this study suggests that investigation of human cardiovascular disease should expand beyond atherosclerotic plaques and left ventricle to gain insight into the pathological remodeling occurring in distant, yet functionally important parts of the heart that may play a role in sustenance and progression of the disease.

## Acknowledgments

We thank Ms Sari Mäki and Ms Teija Kanasuo for their contribution to the immunostainings and multiplex cytokine measurements, respectively.

## Funding

This work was supported by: The Academy of Finland grants 342074 (SLK), 333021 (MUK), 325510 (PT); Clinical Research Fund (EVO) of Turku University Hospital, Turku, Finland (TK); European Research Council (ERC) grant 802825 (MUK); Finnish Association of Cardiothoracic Surgery (KH); Finnish Cardiac Society (KH); Finnish Foundation for Cardiovascular Research (SLK, PRM, MUK, PT, and TK); Finnish Government Research Funding (KH); Finnish Medical Foundation (TK); Ida Montin Foundation (PRM); Instrumentarium Science Foundation (PRM); Saastamoinen Foundation (ES); Sigrid Jusélius Foundation (SLK, MUK, and PT); Orion Research Foundation (SLK, ES, and KH); Yrjö Jahnsson Foundation (SLK, ES)

## Author contributions

Conceptualization: SLK; Formal analysis: SLK, ES, JO, CAB, TÖ, PRM, LH, KK, LZ; Methodology: SLK, KG, TÖ, AT, PKS, PRM, VL, KH, AB, ÅS, EM, MH, MUK, GGC, TK; Investigation: SLK, KG, TÖ, AT, PKS, PRM, VL, KH, AB, ÅS, EM, MH, MUK, GGC, TK; Visualization: SLK, ES, CAB, TÖ, PKS, VL, EM, MH, MUK, GGC, TK; Resources: SLK, KH, HK, JHal, JJ, JG, CAM, MH, JHar, MUK, GGC, PT, TK, MK; Software: SLK, ES, JO, CAB, LH, KK; Funding acquisition: SLK, ES, KH, PRM, MUK, PT, TK; Project administration: SLK; Supervision: SLK, MUK, GGC, PT, TK, MK; Writing – original draft: SLK, ES, KG, TÖ, MH, MUK, TK; Writing – review & editing: SLK, JHal, JHar, GGC, PT, MK

## Competing interests

Authors declare that they have no competing interests.

## Data and materials availability

Data and code available upon request after publication.

## Supplementary Materials

Materials and Methods

Figs. S1 to S10

Tables S1 to S2

References (*88–130*)

## Notes

### Competing Interest Statement

The authors have declared no competing interest.

### Summary of Updates

Major revision. New data added to all sections, all figures/text revised to reflect the update.

## References

1. Wang, L. et al. Single-cell reconstruction of the adult human heart during heart failure and recovery reveals the cellular landscape underlying cardiac function. Nat. Cell Biol. 22, 108–119 (2020).

2. Litviňuková, M. et al. Cells of the adult human heart. Nature 588, 466–472 (2020).

3. Tucker, N. R. et al. Transcriptional and Cellular Diversity of the Human Heart. Circulation 142, 466–482 (2020).

4. Hocker, J. D. et al. Cardiac cell type-specific gene regulatory programs and disease risk association. Sci Adv 7, (2021).

5. Nicin, L. et al. A human cell atlas of the pressure-induced hypertrophic heart. Nature Cardiovascular Research 1, 174–185 (2022).

6. Kuppe, C. et al. Spatial multi-omic map of human myocardial infarction. Nature 608, 766– 777 (2022).

7. Reichart, D. et al. Pathogenic variants damage cell composition and single cell transcription in cardiomyopathies. Science 377, eabo1984 (2022).

8. Koenig, A. L. et al. Single-cell transcriptomics reveals cell-type-specific diversification in human heart failure. Nat Cardiovasc Res 1, 263–280 (2022).

9. Chaffin, M. et al. Single-nucleus profiling of human dilated and hypertrophic cardiomyopathy. Nature 608, 174–180 (2022).

10. 10. Hansen, B. J., Csepe, T. A. & Fedorov, V. V. 28 - Mechanisms of Normal and Dysfunctional Sinoatrial Nodal Excitability and Propagation. in Cardiac Electrophysiology: From Cell to Bedside (Seventh Edition) (eds. Zipes, D. P., Jalife, J. & Stevenson, W. G.) 259–271 (Elsevier, 2018).

11. Wallace, M. J. et al. Genetic Complexity of Sinoatrial Node Dysfunction. Front. Genet. 12, 654925 (2021).

12. Kornej, J., Börschel, C. S., Benjamin, E. J. & Schnabel, R. B. Epidemiology of Atrial Fibrillation in the 21st Century: Novel Methods and New Insights. Circ. Res. 127, 4–20 (2020).

13. Morillo, C. A., Banerjee, A., Perel, P., Wood, D. & Jouven, X. Atrial fibrillation: the current epidemic. J. Geriatr. Cardiol. 14, 195–203 (2017).

14. Berry, C. & Duncker, D. J. Coronary microvascular disease: the next frontier for Cardiovascular Research. Cardiovasc. Res. 116, 737–740 (2020).

15. Vancheri, F., Longo, G., Vancheri, S. & Henein, M. Coronary Microvascular Dysfunction. J. Clin. Med. Res. 9, (2020).

16. Kaski, J.-C., Crea, F., Gersh, B. J. & Camici, P. G. Reappraisal of Ischemic Heart Disease. Circulation 138, 1463–1480 (2018).

17. Godo, S., Suda, A., Takahashi, J., Yasuda, S. & Shimokawa, H. Coronary Microvascular Dysfunction. Arterioscler. Thromb. Vasc. Biol. 41, 1625–1637 (2021).

18. Pepine, C. J. et al. Coronary microvascular reactivity to adenosine predicts adverse outcome in women evaluated for suspected ischemia results from the National Heart, Lung and Blood Institute WISE (Women’s Ischemia Syndrome Evaluation) study. J. Am. Coll. Cardiol. 55, 2825–2832 (2010).

19. Sara, J. D. et al. Prevalence of Coronary Microvascular Dysfunction Among Patients With Chest Pain and Nonobstructive Coronary Artery Disease. JACC Cardiovasc. Interv. 8, 1445–1453 (2015).

20. Hage, C. et al. Association of Coronary Microvascular Dysfunction With Heart Failure Hospitalizations and Mortality in Heart Failure With Preserved Ejection Fraction: A Follow-up in the PROMIS-HFpEF Study. J. Card. Fail. 26, 1016–1021 (2020).

21. Radico, F. et al. Determinants of long-term clinical outcomes in patients with angina but without obstructive coronary artery disease: a systematic review and meta-analysis. Eur. Heart J. 39, 2135–2146 (2018).

22. Lanza, G. A., Crea, F. & Kaski, J. C. Clinical outcomes in patients with primary stable microvascular angina: is the jury still out? Eur Heart J Qual Care Clin Outcomes 5, 283–291 (2019).

23. Camici, P. G. & Crea, F. Coronary microvascular dysfunction. N. Engl. J. Med. 356, 830– 840 (2007).

24. Phan, A., Shufelt, C. & Merz, C. N. B. Persistent chest pain and no obstructive coronary artery disease. JAMA 301, 1468–1474 (2009).

25. Schmauch, E. et al. QClus: Robust and reliable preprocessing method for human heart snRNA-seq. bioRxiv 2022.10.21.513315 (2022) doi:10.1101/2022.10.21.513315.

26. Kalucka, J., de Rooij, L., Goveia, J. & Rohlenova, K. Single-cell transcriptome atlas of murine endothelial cells. Cell (2020).

27. Parmar, K. M. et al. Integration of flow-dependent endothelial phenotypes by Kruppel-like factor 2. J. Clin. Invest. 116, 49–58 (2006).

28. Sangwung, P. et al. KLF2 and KLF4 control endothelial identity and vascular integrity. JCI Insight 2, e91700 (2017).

29. Gimbrone, M. A., Jr & García-Cardeña, G. Endothelial Cell Dysfunction and the Pathobiology of Atherosclerosis. Circ. Res. 118, 620–636 (2016).

30. He, L. et al. NEBULA is a fast negative binomial mixed model for differential or co-expression analysis of large-scale multi-subject single-cell data. Commun Biol 4, 629 (2021).

31. Krämer, A., Green, J., Pollard, J., Jr & Tugendreich, S. Causal analysis approaches in Ingenuity Pathway Analysis. Bioinformatics 30, 523–530 (2014).

32. Sun, C. et al. Functions of exogenous FGF signals in regulation of fibroblast to myofibroblast differentiation and extracellular matrix protein expression. Open Biol. 12, 210356 (2022).

33. Smiljic, S. The clinical significance of endocardial endothelial dysfunction. Medicina 53, 295–302 (2017).

34. Rossjohn, J. et al. T cell antigen receptor recognition of antigen-presenting molecules. Annu. Rev. Immunol. 33, 169–200 (2015).

35. Appay, V. & Rowland-Jones, S. L. RANTES: a versatile and controversial chemokine. Trends Immunol. 22, 83–87 (2001).

36. Sung, M. M. et al. Cardiomyocyte-specific ablation of CD36 accelerates the progression from compensated cardiac hypertrophy to heart failure. Am. J. Physiol. Heart Circ. Physiol. 312, H552–H560 (2017).

37. Dobrzyn, P. et al. Expression of lipogenic genes is upregulated in the heart with exercise training-induced but not pressure overload-induced left ventricular hypertrophy. Am. J. Physiol. Endocrinol. Metab. 304, E1348–58 (2013).

38. Shu, H. et al. The role of CD36 in cardiovascular disease. Cardiovasc. Res. 118, 115–129 (2022).

39. Chandra, M., Miriyala, S. & Panchatcharam, M. PPARγ and Its Role in Cardiovascular Diseases. PPAR Res. 2017, 6404638 (2017).

40. Altara, R. et al. CXCL10 Is a Circulating Inflammatory Marker in Patients with Advanced Heart Failure: a Pilot Study. J. Cardiovasc. Transl. Res. 9, 302–314 (2016).

41. Abbate, A. et al. Interleukin-1 and the Inflammasome as Therapeutic Targets in Cardiovascular Disease. Circ. Res. 126, 1260–1280 (2020).

42. Ridker, P. M. et al. Antiinflammatory Therapy with Canakinumab for Atherosclerotic Disease. N. Engl. J. Med. 377, 1119–1131 (2017).

43. Hickish, T. et al. MABp1 as a novel antibody treatment for advanced colorectal cancer: a randomised, double-blind, placebo-controlled, phase 3 study. Lancet Oncol. 18, 192–201 (2017).

44. Apte, R. S., Chen, D. S. & Ferrara, N. VEGF in Signaling and Disease: Beyond Discovery and Development. Cell 176, 1248–1264 (2019).

45. Gimbrone, M. A., Jr & García-Cardeña, G. Vascular endothelium, hemodynamics, and the pathobiology of atherosclerosis. Cardiovasc. Pathol. 22, 9–15 (2013).

46. Hayes, J. D. & Dinkova-Kostova, A. T. The Nrf2 regulatory network provides an interface between redox and intermediary metabolism. Trends Biochem. Sci. 39, 199–218 (2014).

47. Popov, L.-D. Mitochondrial biogenesis: An update. J. Cell. Mol. Med. 24, 4892–4899 (2020).

48. Dhakshinamoorthy, S., Jain, A. K., Bloom, D. A. & Jaiswal, A. K. Bach1 competes with Nrf2 leading to negative regulation of the antioxidant response element (ARE)-mediated NAD(P)H:quinone oxidoreductase 1 gene expression and induction in response to antioxidants. J. Biol. Chem. 280, 16891–16900 (2005).

49. Nakamura, M. & Sadoshima, J. Mechanisms of physiological and pathological cardiac hypertrophy. Nat. Rev. Cardiol. 15, 387–407 (2018).

50. Dewberry, R., Holden, H., Crossman, D. & Francis, S. Interleukin-1 receptor antagonist expression in human endothelial cells and atherosclerosis. Arterioscler. Thromb. Vasc. Biol. 20, 2394–2400 (2000).

51. Maguire, E. M., Pearce, S. W. A. & Xiao, Q. Foam cell formation: A new target for fighting atherosclerosis and cardiovascular disease. Vascul. Pharmacol. 112, 54–71 (2019).

52. Park, J.-E. et al. A cell atlas of human thymic development defines T cell repertoire formation. Science 367, (2020).

53. Wirka, R. C. et al. Atheroprotective roles of smooth muscle cell phenotypic modulation and the TCF21 disease gene as revealed by single-cell analysis. Nat. Med. 25, 1280–1289 (2019).

54. Zhang, Q. et al. Landscape and Dynamics of Single Immune Cells in Hepatocellular Carcinoma. Cell 179, 829–845.e20 (2019).

55. Zilionis, R. et al. Single-Cell Transcriptomics of Human and Mouse Lung Cancers Reveals Conserved Myeloid Populations across Individuals and Species. Immunity 50, 1317–1334.e10 (2019).

56. Eraslan, G. et al. Single-nucleus cross-tissue molecular reference maps to decipher disease gene function. bioRxiv 2021.07.19.452954 (2021) doi:10.1101/2021.07.19.452954.

57. van den Hoogen, P., et al. Increased circulating IgG levels, myocardial immune cells and IgG deposits support a role for an immune response in pre- and end-stage heart failure. J. Cell. Mol. Med. 23, 7505–7516 (2019).

58. Gerçek, M. et al. Cardiomyocyte Hypertrophy in Arrhythmogenic Cardiomyopathy. Am. J. Pathol. 187, 752–766 (2017).

59. Jaitin, D. A. et al. Lipid-Associated Macrophages Control Metabolic Homeostasis in a Trem2-Dependent Manner. Cell 178, 686–698.e14 (2019).

60. Fatehi Hassanabad, A., Fedak, P. W. M. & Deniset, J. F. Acute Ischemia Alters Human Pericardial Fluid Immune Cell Composition. JACC Basic Transl Sci 6, 765–767 (2021).

61. Ashida, N., Arai, H., Yamasaki, M. & Kita, T. Distinct signaling pathways for MCP-1- dependent integrin activation and chemotaxis. J. Biol. Chem. 276, 16555–16560 (2001).

62. Gibaldi, D. et al. CCL3/Macrophage Inflammatory Protein-1α Is Dually Involved in Parasite Persistence and Induction of a TNF- and IFNγ-Enriched Inflammatory Milieu in Trypanosoma cruzi-Induced Chronic Cardiomyopathy. Front. Immunol. 11, 306 (2020).

63. Luster, A. D., Unkeless, J. C. & Ravetch, J. V. Gamma-interferon transcriptionally regulates an early-response gene containing homology to platelet proteins. Nature 315, 672–676 (1985).

64. Altara, R., Mallat, Z., Booz, G. W. & Zouein, F. A. The CXCL10/CXCR3 Axis and Cardiac Inflammation: Implications for Immunotherapy to Treat Infectious and Noninfectious Diseases of the Heart. J Immunol Res 2016, 4396368 (2016).

65. Douglas, G. et al. A key role for the novel coronary artery disease gene JCAD in atherosclerosis via shear stress mechanotransduction. Cardiovasc. Res. 116, 1863–1874 (2020).

66. Zhao, C. et al. Supervillin promotes tumor angiogenesis in liver cancer. Oncol. Rep. 44, 674–684 (2020).

67. Wang, S. et al. PALMD regulates aortic valve calcification via altered glycolysis and NF- κB-mediated inflammation. J. Biol. Chem. 298, 101887 (2022).

68. Wang, X., Lu, Y., Xie, Y., Shen, J. & Xiang, M. Emerging roles of proteoglycans in cardiac remodeling. Int. J. Cardiol. 278, 192–198 (2019).

69. van der Harst, P. & Verweij, N. Identification of 64 Novel Genetic Loci Provides an Expanded View on the Genetic Architecture of Coronary Artery Disease. Circ. Res. 122, 433–443 (2018).

70. Yan, J. et al. Systematic analysis of binding of transcription factors to noncoding variants. Nature 591, 147–151 (2021).

71. Kheradpour, P. & Kellis, M. Systematic discovery and characterization of regulatory motifs in ENCODE TF binding experiments. Nucleic Acids Res. 42, 2976–2987 (2014).

72. Wang, L., Lin, L., Qi, H., Chen, J. & Grossfeld, P. Endothelial Loss of ETS1 Impairs Coronary Vascular Development and Leads to Ventricular Non-Compaction. Circ. Res. 131, 371– 387 (2022).

73. King, K. R. et al. IRF3 and type I interferons fuel a fatal response to myocardial infarction. Nat. Med. 23, 1481–1487 (2017).

74. Tuomainen, T. & Tavi, P. The role of cardiac energy metabolism in cardiac hypertrophy and failure. Exp. Cell Res. 360, 12–18 (2017).

75. Dinarello, C. A. Overview of the IL-1 family in innate inflammation and acquired immunity. Immunol. Rev. 281, 8–27 (2018).

76. Ridker, P. M. & Rane, M. Interleukin-6 Signaling and Anti-Interleukin-6 Therapeutics in Cardiovascular Disease. Circ. Res. 128, 1728–1746 (2021).

77. Ross, R. Atherosclerosis--an inflammatory disease. N. Engl. J. Med. 340, 115–126 (1999).

78. Ridker, P. M., Cushman, M., Stampfer, M. J., Tracy, R. P. & Hennekens, C. H. Inflammation, aspirin, and the risk of cardiovascular disease in apparently healthy men. N. Engl. J. Med. 336, 973–979 (1997).

79. Ridker, P. M., Hennekens, C. H., Buring, J. E. & Rifai, N. C-reactive protein and other markers of inflammation in the prediction of cardiovascular disease in women. N. Engl. J. Med. 342, 836–843 (2000).

80. Zhang, D., Jiang, S. L., Rzewnicki, D., Samols, D. & Kushner, I. The effect of interleukin- 1 on C-reactive protein expression in Hep3B cells is exerted at the transcriptional level. Biochem. J 310 (Pt 1), 143–148 (1995).

81. 81. C Reactive Protein Coronary Heart Disease Genetics Collaboration (CCGC), et al. Association between C reactive protein and coronary heart disease: mendelian randomisation analysis based on individual participant data. BMJ 342, d548 (2011).

82. Emerging Risk Factors Collaboration, et al. C-reactive protein concentration and risk of coronary heart disease, stroke, and mortality: an individual participant meta-analysis. Lancet 375, 132–140 (2010).

83. Ridker, P. M. et al. Relationship of C-reactive protein reduction to cardiovascular event reduction following treatment with canakinumab: a secondary analysis from the CANTOS randomised controlled trial. Lancet 391, 319–328 (2018).

84. Ridker, P. M. et al. Modulation of the interleukin-6 signalling pathway and incidence rates of atherosclerotic events and all-cause mortality: analyses from the Canakinumab Anti-Inflammatory Thrombosis Outcomes Study (CANTOS). Eur. Heart J. 39, 3499–3507 (2018).

85. Ridker, P. M. et al. IL-6 inhibition with ziltivekimab in patients at high atherosclerotic risk (RESCUE): a double-blind, randomised, placebo-controlled, phase 2 trial. Lancet 397, 2060–2069 (2021).

86. Kleveland, O. et al. Effect of a single dose of the interleukin-6 receptor antagonist tocilizumab on inflammation and troponin T release in patients with non-ST-elevation myocardial infarction: a double-blind, randomized, placebo-controlled phase 2 trial. Eur. Heart J. 37, 2406– 2413 (2016).

87. Broch, K. et al. Randomized Trial of Interleukin-6 Receptor Inhibition in Patients With Acute ST-Segment Elevation Myocardial Infarction. J. Am. Coll. Cardiol. 77, 1845–1855 (2021).

88. Kuosmanen, S. M. et al. MicroRNA profiling of pericardial fluid samples from patients with heart failure. PLoS One 10, e0119646 (2015).

89. Sousa-Uva, M. et al. 2018 ESC/EACTS Guidelines on myocardial revascularization. Eur. J. Cardiothorac. Surg. 55, 4–90 (2019).

90. 90. Lawton, J. S. et al. 2021 ACC/AHA/SCAI Guideline for Coronary Artery Revascularization: A Report of the American College of Cardiology/American Heart Association Joint Committee on Clinical Practice Guidelines. Circulation 145, e18–e114 (2022).

91. Ponikowski, P. et al. 2016 ESC Guidelines for the diagnosis and treatment of acute and chronic heart failure: The Task Force for the diagnosis and treatment of acute and chronic heart failure of the European Society of Cardiology (ESC)Developed with the special contribution of the Heart Failure Association (HFA) of the ESC. Eur. Heart J. 37, 2129–2200 (2016).

92. Hollenberg, S. M. et al. 2019 ACC Expert Consensus Decision Pathway on Risk Assessment, Management, and Clinical Trajectory of Patients Hospitalized With Heart Failure: A Report of the American College of Cardiology Solution Set Oversight Committee. J. Am. Coll. Cardiol. 74, 1966–2011 (2019).

93. Wolf, F. A., Angerer, P. & Theis, F. J. SCANPY: large-scale single-cell gene expression data analysis. Genome Biol. 19, 15 (2018).

94. Wolock, S. L., Lopez, R. & Klein, A. M. Scrublet: Computational Identification of Cell Doublets in Single-Cell Transcriptomic Data. Cell Syst 8, 281–291.e9 (2019).

95. Korsunsky, I. et al. Fast, sensitive and accurate integration of single-cell data with Harmony. Nat. Methods 16, 1289–1296 (2019).

96. McInnes, L., Healy, J. & Melville, J. UMAP: Uniform Manifold Approximation and Projection for Dimension Reduction. arXiv [stat.ML] (2018).

97. Becht, E. et al. Dimensionality reduction for visualizing single-cell data using UMAP. Nat. Biotechnol. (2018) doi:10.1038/nbt.4314.

98. Traag, V. A., Waltman, L. & van Eck, N. J. From Louvain to Leiden: guaranteeing well-connected communities. Sci. Rep. 9, 5233 (2019).

99. Kuleshov, M. V. et al. Enrichr: a comprehensive gene set enrichment analysis web server 2016 update. Nucleic Acids Res. 44, W90–7 (2016).

100. Fang, Z. GSEApy: Gene Set Enrichment Analysis in Python. (Github).

101. Benjamini, Y. & Hochberg, Y. Controlling the False Discovery Rate: A Practical and Powerful Approach to Multiple Testing. J. R. Stat. Soc. Series B Stat. Methodol. 57, 289–300 (1995).

102. Parmar, K. M. et al. Integration of flow-dependent endothelial phenotypes by Kruppel-like factor 2. J. Clin. Invest. 116, 49–58 (2006).

103. Ewels, P. A. et al. The nf-core framework for community-curated bioinformatics pipelines. Nat. Biotechnol. 38, 276–278 (2020).

104. Dobin, A. et al. STAR: ultrafast universal RNA-seq aligner. Bioinformatics 29, 15–21 (2013).

105. Patro, R., Duggal, G., Love, M. I., Irizarry, R. A. & Kingsford, C. Salmon provides fast and bias-aware quantification of transcript expression. Nat. Methods 14, 417–419 (2017).

106. Love, M. I., Huber, W. & Anders, S. Moderated estimation of fold change and dispersion for RNA-seq data with DESeq2. Genome Biol. 15, 550 (2014).

107. ENCODE Project Consortium. An integrated encyclopedia of DNA elements in the human genome. Nature 489, 57–74 (2012).

108. Heinz, S. et al. Simple combinations of lineage-determining transcription factors prime cis-regulatory elements required for macrophage and B cell identities. Mol. Cell 38, 576–589 (2010).

109. Ritchie, M. E. et al. limma powers differential expression analyses for RNA-sequencing and microarray studies. Nucleic Acids Res. 43, e47 (2015).

110. Sha, Y., Phan, J. H. & Wang, M. D. Effect of low-expression gene filtering on detection of differentially expressed genes in RNA-seq data. Conf. Proc. IEEE Eng. Med. Biol. Soc. 2015, 6461–6464 (2015).

111. Howe, K. L., et al. Ensembl 2021. Nucleic Acids Res. 49, D884–D891 (2021).

112. Palla, G. et al. Squidpy: a scalable framework for spatial omics analysis. Nat. Methods 19, 171–178 (2022).

113. Chistiakov, D. A., Killingsworth, M. C., Myasoedova, V. A., Orekhov, A. N. & Bobryshev, Y. V. CD68/macrosialin: not just a histochemical marker. Lab. Invest. 97, 4–13 (2017).

114. Betjes, M. G., Haks, M. C., Tuk, C. W. & Beelen, R. H. Monoclonal antibody EBM11 (anti-CD68) discriminates between dendritic cells and macrophages after short-term culture. Immunobiology 183, 79–87 (1991).

115. Murray, P. J. & Wynn, T. A. Protective and pathogenic functions of macrophage subsets. Nat. Rev. Immunol. 11, 723–737 (2011).

116. Mountjoy, E. et al. An open approach to systematically prioritize causal variants and genes at all published human GWAS trait-associated loci. Nat. Genet. 53, 1527–1533 (2021).

117. Nelson, C. P. et al. Association analyses based on false discovery rate implicate new loci for coronary artery disease. Nat. Genet. 49, 1385–1391 (2017).

118. Malik, R. et al. Multiancestry genome-wide association study of 520,000 subjects identifies 32 loci associated with stroke and stroke subtypes. Nat. Genet. 50, 524–537 (2018).

119. Kichaev, G. et al. Leveraging Polygenic Functional Enrichment to Improve GWAS Power. Am. J. Hum. Genet. 104, 65–75 (2019).

120. Matsunaga, H., et al. Transethnic Meta-Analysis of Genome-Wide Association Studies Identifies Three New Loci and Characterizes Population-Specific Differences for Coronary Artery Disease. Circ Genom Precis Med 13, e002670 (2020).

121. Koyama, S. et al. Population-specific and trans-ancestry genome-wide analyses identify distinct and shared genetic risk loci for coronary artery disease. Nat. Genet. 52, 1169–1177 (2020).

122. Hartiala, J. A. et al. Genome-wide analysis identifies novel susceptibility loci for myocardial infarction. Eur. Heart J. 42, 919–933 (2021).

123. Kurki, M. I. et al. FinnGen: Unique genetic insights from combining isolated population and national health register data. bioRxiv (2022) doi:10.1101/2022.03.03.22271360.

124. Zhou, W. et al. Efficiently controlling for case-control imbalance and sample relatedness in large-scale genetic association studies. Nat. Genet. 50, 1335–1341 (2018).

125. Aragam, K. G. et al. Discovery and systematic characterization of risk variants and genes for coronary artery disease in over a million participants. bioRxiv (2021) doi:10.1101/2021.05.24.21257377.

126. Li, L. et al. Transcriptome-wide association study of coronary artery disease identifies novel susceptibility genes. Basic Res. Cardiol. 117, 6 (2022).

127. Erdmann, J., Kessler, T., Munoz Venegas, L. & Schunkert, H. A decade of genome-wide association studies for coronary artery disease: the challenges ahead. Cardiovasc. Res. 114, 1241– 1257 (2018).

128. Muerdter, F. et al. Resolving systematic errors in widely used enhancer activity assays in human cells. Nat. Methods 15, 141–149 (2018).

129. Toropainen, A. et al. Functional noncoding SNPs in human endothelial cells fine-map vascular trait associations. Genome Res. 32, 409–424 (2022).

130. Örd, T. et al. Single-Cell Epigenomics and Functional Fine-Mapping of Atherosclerosis GWAS Loci. Circ. Res. 129, 240–258 (2021).

