## Supplemental Files for "Cardiovascular disease causes proinflammatory microvascular changes in the human right atrium"

### Materials and Methods

#### Sample selection

Human right atrial appendage tissue biopsies were harvested as a part of the ongoing prospective PERIHEART and CAREBANK studies (ClinicalTrials.gov Identifier: NCT03444259). Ethical Committees of the Hospital Districts of Southwest Finland and Northern Savo approved the protocol and study complies with the Declaration of Helsinki as revised in 2002. The CAREBANK study (**Table 1a**) has enrolled patients undergoing open-heart cardiac surgery (coronary bypass surgery, operations for valvular heart disease and ascending aorta) since February 2016 at Turku University Hospital and the PERIHEART study (**Table 1b**) since 2012. All patients gave their written informed consent prior to surgery. Samples were taken from the right atrial appendage, from the anatomical site of cardiopulmonary bypass venous cannulation. They were always obtained in the beginning of the operation immediately before patient was coupled to cardiopulmonary bypass to avoid the effects of cardiopulmonary bypass time or technical surgery type on the sample quality. Pericardial fluid was collected in the beginning of the surgery by aspirating to a syringe. Tissue samples were snap-frozen immediately upon collection and stored at -80°C. Pericardial fluid samples were processed as described in Kuosmanen *et al.* (*88*) and all tubes were snap-frozen and stored as tissue samples. Clinical data collection was performed prospectively in an electronic case report form by a trained research nurse with structured questionnaires. In the CAREBANK samples, patients with ischemic heart disease had an indication for coronary revascularization as appropriate by the current guidelines (*89,90*). Ischemic heart failure was defined as reduced left ventricular ejection fraction below 50% in echocardiography in addition to above criteria for revascularization (*91,92*). Non-ischemic heart failure was defined as reduced left ventricular ejection fraction below 50% without coronary artery disease in preoperative coronary angiogram. In the PERIHEART study, control individuals were operated due to mitral valve regurgitation and had no other significant cardiovascular disease. Stable coronary artery disease group (stable CAD) consisted of patients with significant coronary artery disease who underwent elective Coronary Artery Bypass Grafting (CABG) surgery and had no other significant cardiovascular disease (*89,90*). Patients with acute myocardial infarction (acute MI) had preoperative myocardial infarction and were operated with CABG during the same hospital stay. Patients with remote myocardial infarction (remote MI) had suffered myocardial infarction in the past but not prior to CABG surgery.

**Table S2a.** CAREBANK samples

| | Control | IHD | IHF | NIHF |
|---|---|---|---|---|
| N | 6 | 11 | 11 | 3 |
| Gender | 4M 2F | 9M 2F | 10M 1F | 2M 1F |
| Age (years) * | 67± 4 | 68 ± 8 | 66 ± 13 | 72 ± 6 |
| BMI (kg/m2) * | 29 ± 4 | 30 ± 4 | 28 ± 5 | 29 ± 4 |
| Smoker | 0 | 0 | 4 | 0 |
| Atrial fibrillation | 0 | 0 | 2 | 2 |
| Treatment for hypertension | 4 | 9 | 9 | 3 |
| Treatment for diabetes | 2 | 4 | 6 | 0 |
| Treatment for Hypercholesterolemia | 3 | 10 | 11 | 1 |
| Heart failure | 0 | 0 | 5 | 3 |
| Prior Stroke / TIA | 1 | 0 | 1 | 0 |
| Previous myocardial infarction | 0 | 0 | 4 | 0 |
| Previous PCI | 0 | 3 | 2 | 0 |
| Previous CABG | 0 | 0 | 0 | 0 |

| | | | | |
|---|---|---|---|---|
| Number of stenosis in the coronary arteries | 0 | 2 | 2 | 0 |
| LVEF* | 64 ± 4 | 64 ± 7 | 38 ± 7 | 41 ± 9 |
| Betablocker | 2 | 10 | 9 | 3 |
| Calciumblocker | 0 | 3 | 4 | 2 |
| ACE-inhibitor | 5 | 7 | 10 | 3 |
| Warfarin | 0 | 0 | 3 | 0 |
| NOAC | 0 | 1 | 1 | 1 |
| ASA | 4 | 9 | 7 | 0 |
| ADPB | 0 | 1 | 1 | 0 |
| Statin | 3 | 10 | 11 | 1 |
| Insuline | 0 | 3 | 4 | 0 |

*Mean ± Standard deviation*

**Table S2b.** PERIHEART samples

| | Control | stable CAD | acute MI | remote MI |
|---|---|---|---|---|
| N | 4 | 5 | 4 | 5 |
| Gender | all male | all male | all male | all male |
| Age (years) * | 61±7 | 68 ±4 | 66 ±6 | 72 ±6 |
| BMI (kg/m2) * | 26 ±4 | 26 ±3 | 25 ±2 | 28 ±3 |
| Smoker | 0 | 0 | 2 | 2 |
| Atrial fibrillation | 0 | 0 | 0 | 0 |
| Treatment for hypertension | 3 | 4 | 3 | 3 |
| Treatment for diabetes | 0 | 1 | 1 | 2 |

| | | | | |
|---|---|---|---|---|
| Treatment for Hypercholesterolemia | 1 | 3 | 4 | 3 |
| Heart failure | 0 | 0 | 2 | 1 |
| Prior Stroke / TIA | 0 | 0 | 0 | 0 |
| Previous myocardial infarction | 0 | 0 | 4 | 5 |
| Previous PCI | 0 | 0 | 1 | 3 |
| Previous CABG | 0 | 0 | 0 | 0 |
| Number of stenosis in the coronary arteries | 0 | 3 | 3 | 3 |
| LVEF * | 75 ±4 | 61 ±7 | 49 ±13 | 48 ±9 |
| Betablocker | 2 | 2 | 3 | 5 |
| Calciumblocker | 0 | 0 | 2 | 0 |
| ACE-inhibitor | 1 | 1 | 2 | 4 |
| Warfarin | 0 | 0 | 0 | 2 |
| NOAC | 0 | 0 | 0 | 0 |
| ASA | 0 | 4 | 3 | 4 |
| ADPB | 0 | 0 | 1 | 1 |
| Statin | 1 | 4 | 3 | 5 |
| Insuline | 0 | 0 | 0 | 0 |

**Mean ± Standard deviation*

Histochemistry

Tissues were embedded in Optimal cutting temperature (OCT) compound and stored at -80ºC. OCT-embedded tissue blocks were sectioned at a thickness of 10 microns. Frozen tissue sections were air-dried at room temperature and then fixed in 4% paraformaldehyde in PBS solution for 10 min. Fixed tissue sections were stained with Hematoxylin (Vector Laboratories, Burlingame, CA) for 10 min. After several washes in distilled water, sections were then stained with 0.5% Eosin (Sigma-Aldrich, St. Louis, MO) for 10 s followed by several washes in distilled water. Sections were then dehydrated by sequentially dipping 10 times in 50%, 70%, 95% and 100% ethanol respectively. Slides were completely air-dried and then cleared by dipping in xylene for 10s. Stained tissue sections were mounted using EcoMount mounting medium (Biocare Medical, Pacheco, CA) and images were captured with a Nikon DX-DB digital camera using a Nikon Digital Sight DS-U1 microscope.

Isolation of nuclei from ex vivo cardiac tissue and pericardial fluid

Pericardial fluid was processed using a three-step centrifugation protocol as described in Kuosmanen *et al*. (*88*), and the cell pellets were flash-frozen immediately after collection. For nuclei isolation, the frozen pellet was thawed for 3 min in a water bath at 37ºC, spun down and treated with 0.5 ml ACK lysing buffer (Gibco, Thermo Fisher Scientific, Inc) for 5 min at RT. Cells were centrifuged for 5 min, at 300 x g at 4ºC, the supernatant was removed, and the cleared pellet was lysed in 0.1 ml lysis buffer (10 mM Tris-HCl (pH 7.4), 10 mM NaCl, 3 mM MgCl, 0.1% Tween 20, 0.1% NP40, 1% BSA, 1 mM DDT, 1 U/µl Rnase inhibitors (TaKaRa)). Lysis progression was monitored under the microscope and stopped as soon as 95-98% of cells were lysed with the addition of 1 ml ice-cold wash buffer (10 mM Tris-HCl (pH 7.4), 10 mM NaCl, 3 mM MgCl, 0.1% Tween 20, 1% BSA, 1 mM DDT, 1 U/ul Rnase inhibitors (TaKaRa)). Nuclei were spun down for 5 min, at 500 x g, 4ºC, resuspended in 0.04% BSA in PBS, stained with Trypan Blue, counted on a hemocytometer, and adjusted to a concentration of 1000 nuclei/µl.

For tissue biopsies in OCT, surrounding OCT was removed with a scalpel and the tissue was washed three times with ice-cold 0.04% BSA in PBS. For all tissue biopsies, the tissue was transferred into a tube containing 1 ml of ice-cold lysis buffer (0.32 M Sucrose, 5 mM $CaCl_2$, 3mM MgAc, 2.0 mM EDTA, 0.5mM EGTA, 10 mM Tris-HCl (pH 8.0), 1 mM DDT, EDTA-free protease inhibitor cocktail (Roche, Complete tablet), 1 U/µl RNase inhibitors (TaKaRa)), cut into 2-3 mm pieces with scissors, transferred to an ice-cold glass dounce tissue grinder and after 5 min of incubation on ice, stroked with a "Loose" and a "Tight" pestle, 15 times each. The solution was passed through a 40-micron strainer and centrifuged at 1000 x g for 8 min at 4°C on a swinging bucket rotor. The supernatant was removed, and the nuclear pellet was resuspended in 1 ml of nuclei suspension buffer (0.04% BSA in PBS with RNase inhibitors) and centrifuged at 1000 x g for 8 min at 4°C on a swinging bucket rotor. After removal of the supernatant, nuclear pellet was resuspended in the remaining nuclei suspension buffer. The quality of the nuclear suspension was estimated under the microscope following staining with Trypan Blue and nuclei concentration was adjusted to 1000 nuclei / µl.

Droplet-based snRNA-seq

Immediately after nuclei isolation, single-nuclei RNA (snRNA) libraries were processed using the droplet-based RNA sequencing technology. Briefly, 5000-7000 nuclei were profiled per sample using the Chromium Single Cell 3' RNA reagent kit v3 according to the 10X Genomics protocol. The generated cDNA libraries were indexed, pooled, and sequenced using the NovaSeq 6000 S2 system and reagent kits (100 cycles) (Illumina, Inc).

Bioinformatics preprocessing

The samples were processed using Cell Ranger v6.0 (10x Genomics) and the data were analyzed using Scanpy v1.7 (*93*) on python v3.9. To remove empty droplets and highly contaminated cells, we used a specific method, that we developed especially for cardiac tissue, QClus (*25*). Briefly,

QClus establishes contamination-related quality metrics, such as splicing fraction, nuclear genes expression, mitochondrial fraction, in addition to cardiomyocyte and non-cardiomyocyte gene enrichments. These metrics are used to cluster and filter the nuclei, resulting in cleaner data. The QClus pipeline includes doublet filtering with Scrublet (*94*). After QClus filtering, all samples raw counts were merged, normalized, and then transformed in log values (ln (counts per 10000) + 1). The resulting normalized matrix was used for visualization of gene expression. Correlations for gene and read numbers in heart tissue and pericardial fluid are shown below together with mitocohndrial quality metrics for heart tissue.

Heart :

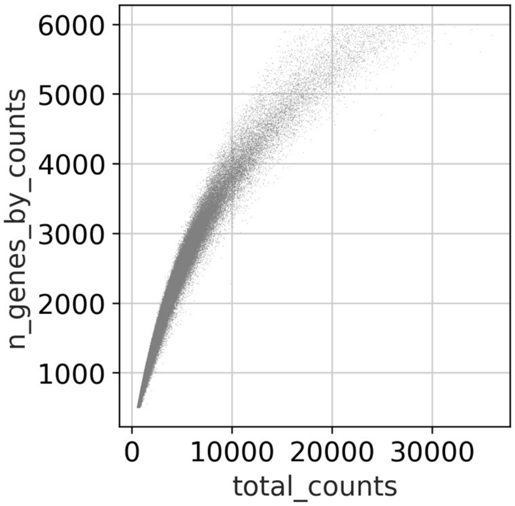


Pericardial fluid :

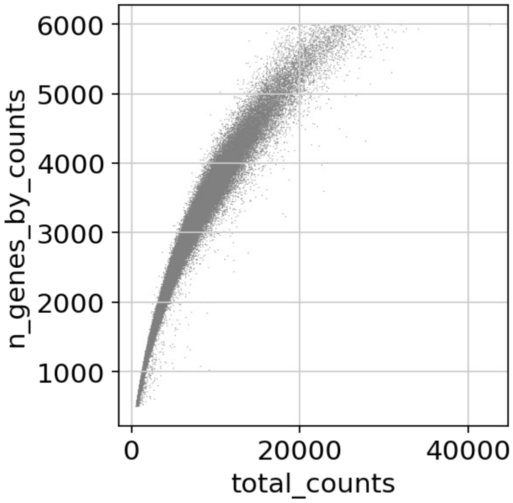


Heart tissue:

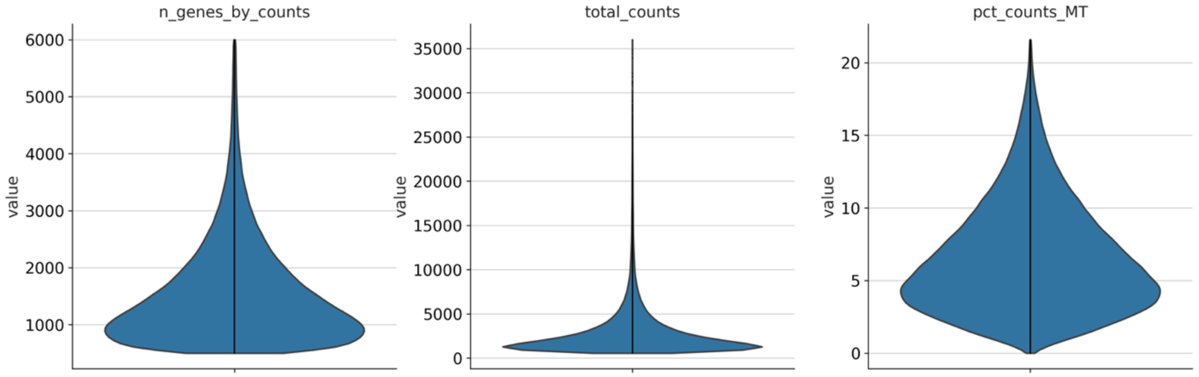

Expression correction, sample heterogeneity correction, embedding and clustering

Genes that were expressed in less than 10 cells (across all samples) were filtered out. Variability filtering of genes was also applied (minimum mean of 0.0125, maximum mean of 3 and minimum normalized dispersion of 0.5). Linear regression was used to correct for the number of counts per cell and mitochondrial percentage. Subsequent steps were standard scaling, Principal Component Analysis (PCA) and Harmony batch correction (*95*) at the sample level. Afterwards, a 10 nearest neighbor graph was constructed, based on the top 40 PCs, which was used for UMAP embedding (*96,97*). Clusters were found using the Leiden algorithm (*98*) at resolution 1.

Cell type identification

Clusters were annotated through cell-type specific marker expression and correlation. Annotations were additionally confirmed through gene set enrichment using Enrichr (*99*) (from gseapy (*100*) wrapper), from the first 500 significantly differentially expressed genes (Wilcoxon ranked sum, Benjamini-Hochberg FDR < 0.05). At the cell-typing step and further subtyping, when a Leiden cluster (*98*) showed high level expression of all markers from two distinct cell-types, it was removed as a doublet cluster. Cell-type specific scores were calculated by using the *score_genes* of scanpy (*93*) with marker genes from these cell-types.

Cell subtyping

Seven main sets of cells were established by combining 12 discovered cell-types, namely cardiomyocytes, endocardial endothelial cells, fibroblasts, mesothelial cells, vascular cells (containing pericytes, smooth muscle cells, and vascular endothelial cells) immune cells (containing lymphocytes and macrophages), and neuro-adipo cells (containing Schwann cells, adipocytes, and neurons). Subclusters were found through the reprocessing of each set, by regression, scaling and PCA steps. Batch correction was also performed again at the sample level, in addition with k nearest neighbors' (knn) graph and UMAP plot. Clusters were discovered with Leiden clustering (resolution 1) (*98*). For each set, small clusters were found to be enriched for markers specific for other cell types and were thus removed as doublets.

Differential expression analysis

Apart from cluster-specific marker identification, differential gene expression analysis was performed using the NEBULA (*30*) R package. Mitochondrial percentage, patient age and patient sex were used as additive confounding covariates in the model. As advised by the documentation, the total number of counts for each cell was imputed as the scaling factor. The LN method was used when the number of cells exceeded 1000, and the HL method when not. Benjamini-Hochberg method (*101*) was used for FDR correction.

RNA-seq

*Disturbed flow*

Human umbilical vein endothelial cells (HUVECs) were isolated and cultured as previously described (*102*). Cells were then exposed to disturbed, non-laminar flow or left under static (no flow) conditions for 24h. At the end of the experiments, cells were lysed, and mRNA was isolated and used for NGS library preparation and sequencing at the Broad Institute.

Fastq files were processed and reads aligned using nf-core RNA-seq pipeline (*103*). Briefly, after QC filtering and trimming, the pipeline used STAR (*104*) to align the reads to the hg38 Genome, and Salmon (*105*) to create a count matrix. The resulting length scaled count matrix was used as an input to Deseq2 (*106*) for differential expression analysis.

*IL-1β time course analysis*

Human aortic endothelial cells (HAEC)(LONZA) were maintained in endothelial cell growth medium (EGM; basal medium with SingleQuots supplements CC-4133; Lonza) supplemented with 10% fetal bovine serum (FBS; GIBCO) on T-75 cell culture flasks coated with 10 g/ml fibronectin (Sigma, St Louis, MO, USA) and 0.05% gelatin at 37°C in a humidified atmosphere at 5% CO2 and used at passage 4 to 6. Prior to IL-1β stimulation, HAEC medium was changed with fresh endothelial cell growth medium (EGM; basal medium; Lonza) supplemented with 1% fetal bovine serum (FBS; GIBCO). HAEC were then stimulated with 10 ng/ml of IL-1β (PHC0814, GIBCO) at 2, 8, and 23h time points. RNA was extracted using RNeasy Mini Kit (74104, QIAGEN) according to manufacturer protocol. Extracted RNA was sent to the Sequencing Service GeneCore Sequencing Facility (EMBL, www.genecore.embl.de) for NGS library preparation using stranded rRNA-depleted RNA-Seq protocol.

The sequencing reads obtained were trimmed to 3' A-stretches originating from the library preparation and poor-quality reads were filtered out (minimum 97% of bp over quality cutoff 10). Remaining reads were aligned to GRCh37hg19 reference genome using STAR v2.5.4b (*104*) with ENCODE recommended options for long RNA-seq analysis (*107*). Reads located on exonic locations were quantified using HOMER v4.9 (*108*) 'analyzeRepeats' routine with the settings '-condense genes -count exons'. Differential expression was calculated using limma version 3.46.0 (*109*) after filtering of low expressed transcripts (CPM >0.5 in at least 2 samples) to improve sensitivity and precision of the analysis (*110*).

Ingenuity Pathway Analysis

The affected 'Diseases and biological functions', 'Canonical pathways' and 'Upstream regulators' were studied using the 'Core analysis' and 'Comparison analysis' of the Ingenuity Pathway Analysis (IPA, QIAGEN Inc., https://digitalinsights.qiagen.com/IPA) (*31*) based on the differentially expressed genes for each cell type / cell subtype. The data tables can be found from the website (**website**). Pathway overlaps were explored using Venny (https://bioinfogp.cnb.csic.es/tools/venny/index.html).

Resolve molecular cartography

Fresh-frozen tissue was sectioned and transferred to capture areas on spatial transcriptomics slides by Resolve Bioscience and processed using their Molecular Cartography™ spatial transcriptomics. One hundred genes were measured at a time. For each gene, all transcripts for ENSEMBL (*111*) were utilized to ensure that they were all counted. Samples were primed and hybridized with all probes which were then fluorescently tagged. Regions of interest were imaged and decolorized to remove fluorescent signals. Development of color, imaging and decolorization steps were repeated several times in cycles to create a combination of fluorescence, which was decoded to retrieve the gene information from the images. The resulting data were analyzed using a tailored method in Python to overlap the spatial expression of genes of interest from specific ROIs. We created a grid of 80px wide hexagons, covering the whole slide, and counted signals within each, simulating small, high-resolution spots. We used the Scanpy (*93*) and Squidpy (*112*) methods to process the resulting data. The samples were merged into one matrix. As spots contained few counts, and only 100 genes were characterized, the data were not normalized to prevent artifacts. The counts were logarithmized (using log1p function of Scanpy) and linear regression was performed to eliminate the effect of global expression levels between spots (number of counts and number of genes) and the data were scaled (standard scaling, max value of 10). Afterwards, PCA was performed, as well as sample-level batch correction, using Harmony (*95*). Then, a knn network was constructed for the creation of a UMAP embedding (*97*), using the top 15 PCs. Clusters were discovered using the Leiden algorithm (*98*). For detailed plotting, each count was plotted as a dot of a particular color, corresponding to a gene, at the exact location where it was associated in the original data, resulting from the experiment.

Visium spatial transcriptomics

One heart failure and one control sample were processed, and four technical replicates were measured for each sample. Thin (10µm) sections of OCT-embedded tissue were cryosectioned at -20°C on a cryostat, mounted on tissue optimization (TO) and gene expression (GEX) Visium slides and stored at -80°C for up to one week. TO slides were used to determine the optimum tissue permeabilization time following the protocol of the manufacturer (Visium Spatial TO Slide and Reagent kit, 10x Genomics, Pleasanton, CA). Briefly, sections were methanol-fixed, stained with hematoxylin and eosin (H&E stain), and imaged on a Leica DMi8 microscope. Next, a permeabilization time course was performed, followed by steps of fluorescent cDNA synthesis, tissue removal, and fluorescent imaging. GEX slides were similarly fixed, H&E stained, and imaged on a Leica DMi8 microscope. Next, the tissue was permeabilized according to the TO-determined permeabilization time and four cDNA libraries were prepared according to the manufacturer protocol (Visium Spatial GEX Slide and Reagent kit, 10x Genomics, Pleasanton, CA). Each set of 4 libraries was pooled and sequenced using the NextSeq 500 system (150 cycles) (Illumina, Inc).

Samples were processed using Scanpy 1.9.1 (*93*). All sample data matrices were merged into one matrix which was then processed. Counts were normalized (total count of 10,000 per capture area) and logarithmized (using log1p function of Scanpy). For dimension reduction purposes, the raw normalized matrix was further processed: genes that were not characterized as highly variable enough were filtered out (minimum mean of 0.0125, maximum mean of 3 and minimum normalized dispersion of 0.5), and linear regression was performed to eliminate the effect of covariates (total_counts, and mitochondrial genes percentage). The data were then scaled (standard scaling, max value of 10). Afterwards, PCA was performed, as well as sample-level batch correction, using Harmony (*95*). Then, a knn network was constructed for the creation of a UMAP embedding (*97*). Clusters were discovered using the Leiden algorithm (*98*). Cell-type-specific scores were calculated by using the *score_genes* of scanpy with marker genes of these cell-types, originating from the snRNA-seq data.

Cytokine screening from serum samples

Patient samples were selected from CAREBANK study (ClinicalTrials.gov Identifier: NCT03444259). Serum samples were collected preoperatively after at least 8h of fasting by routine sampling of the certified hospital laboratory. The serum samples were stored at -70°C until analysis. Serum cytokine screening was done using the Bio-Plex Pro Human Cytokine Screening Panel, 48-Plex (cat#: 12007283, Bio-rad) according to the instructions of the manufacturer.

Immunohistochemistry

For immunostaining, OCT blocks were sectioned at a thickness of 5 micrometers. Standard immunostaining protocols were used. Anti-CD68 antibody (prediluted, cat#: ab845, Abcam) was used as a common macrophage marker, anti-TREM2 antibody (clone: 237920, Novus Biologicals) for identifying LAMs, and BODIPY™ 493/503 (4,4-Difluoro-1,3,5,7,8-Pentamethyl-4-Bora-3a,4a-Diaza-s-Indacene; cat#: D3922, ThermoFisher) for staining lipids (*113-115*). Stained slides were scanned using Panoramic 250 digital slide scanner. Criteria for macrophages included morphology and granular inclusions in the cytoplasm. For each subject the same sized randomly selected areas from pericardium were evaluated. Images were analyzed using ImageJ2/Fuji. Number of CD68 positive macrophages were divided by total nuclei count.

Gene expression modules

We calculated gene expression modules for each major cell class using the scmodule package (manuscript in preparation). Briefly, the method first adjusts the gene expression matrix **X** by performing zero-phase component analysis (ZCA) to remove cell-cell correlation structure, which biases the recovery of gene-gene modules. We calculate the SVD of **X** as $\mathbf{USV^T}$ and use it to approximate the gene-gene correlation matrix of the ZCA adjusted expression matrix as $\mathbf{C_{ZCA}} \sim \mathbf{VS^PV^T}$ at multiple resolutions **P** in [0,0.5,1]. For each matrix, we z-score the estimated correlations controlling for sparsity of each gene (number of zeros in the expression matrix) and build a gene-gene graph with correlations with $z > 4.5$. Finally, we detect modules of genes by multi-graph Leiden clustering (*98*) on the ensemble of graphs (with leiden resolution=2).

GWAS linked genes

Published gene prioritization results were collected across multiple different prioritization approaches and GWAS studies. The OpenTargets Genetics Portal (*116*) (data release: June 2021) was used to obtain data for the GWAS studies of CAD, myocardial infarction, and stroke (*69,117-122*) (FinnGen (*123*) and UK Biobank (*124*) data via the OpenTargets study accessions FINNGEN_R5_I9_CORATHER, SAIGE_411, SAIGE_411_2 and SAIGE_411_4). For OpenTargets, the Locus2Gene algorithm prioritized 238 genes, eQTL colocalization 164 genes, and 'nearest gene' 387 genes.
From the recent CAD GWAS by Aragam *et al.* (*125*), the per-association overall top-prioritized genes contributed 186 genes, PoPS prioritization 386 genes, 'nearest gene' 216 genes, and, for the GWAS 1% FDR threshold associations, 'nearest gene' contributed 716 genes.
The TWAS of CAD by Li *et al.* (*126*) prioritized 114 genes and the CAD GWAS review by Erdmann *et al.* (*127*) listed 373 genes at CAD loci.

Allelic activity reporter assay

STARR-Seq massively parallel reporter assay (*128*) was used to compare the transcriptional activity of SNP alleles in teloHAEC human immortalized aortic endothelial cells (ATCC) and in primary human aortic smooth muscle cells (Thermo Scientific). The library generation, quality control and analysis has been described in detail in Toropainen *et al.* (*129*) and Örd *et al.* (*130*), respectively.

CRISPR-RNP transfection

The human endothelial cells, TeloHAECs were maintained in Vascular Cell Basal Medium (ATCC PCS-100-030), supplemented with Vascular Endothelial Cell Growth Kit-VEGF (ATCC PCS-100-041) and 100 I.U./ml penicillin- 100 (µg/ml) streptomycin. The cells were grown to 70-90% confluency before the experiment. The CRISPR deletion experiment was performed as per standard protocol provided by IDT. Briefly, Alt-R CRISPR-Cas9 crRNA/gRNA, and tracrRNA were dissolved in duplex buffer, and Alt-R CRISPR-Cas9 Electroporation Enhancer (EE) in IDTE, pH 8.0 to a final concentration of 100 µM. The crRNA:tracrRNA duplexes were formed by mixing two oligos at equimolar concentrations in a sterile microcentrifuge, then heated at 95°C for 5 mins. For each electroporation reaction, 36µM of Alt-R S.p. HiFi Cas9 enzyme and crRNA:tracrRNA duplexes were incubated together to form the ribonucleoprotein (RNP) complexes following the manufacturer guidelines. The cells were then washed, trypsinized, and re-washed with warm PBS before preparing the working cell stock in resuspension buffer R (Thermo Fisher Sci. Cat# MPK1096). 50 000 cells were used per 10µl of reaction volume, and three reactions were performed for each well of a 12-well plate. Finally, cells, specific RNP complex, and 10.8uM of Alt-R Cas9 electroporation enhancer were mixed before loading into 10µl Neon Transfection System kit. The electroporation was performed using voltage: 1350V, width: 30ms, pulses: 1 pulse. The newly transfected cells were inoculated immediately to pre-warmed complete cell culture media and allowed to grow for 48h before collection. The genomic DNA and RNA were collected from same sample using Quick-DNA/RNA Miniprep Plus Kit (Zymo, Cat D7003). To confirm the

CRISPR-RNP deletion of the specific target regions, we performed PCR followed by agarose gel electrophoresis using genomic DNA and target specific primer pairs (JCAD1_FWD: GCACTTCCTCCTGCCATAAA and JCAD1_REV: ACACCCAACATCCCTGTATTC; JCAD2_FWD: CCTCTTTGCCTACTTCCTCTTAC and JCAD2_REV: GTGGAACCCTCATTACCTCATC. Then, we performed qPCR using ΔΔCt method to measure the relative gene expression of JCAD/KIAA1462 among collected RNA samples using GAPDH as the house keeping gene to normalize between control and treatments. We used the following primer pairs for qPCR: FH2_KIAA1462: AACAATGACTTAAAGCCCAG and BH2_KIAA1462: ACTGAGGTCATTTGTTTGTC; FH1_GAPDH: TCGGAGTCAACGGATTTG and BH1_GAPDH: CAACAATATCCACTTTACCAGAG.

CRISPRi: Plasmid transfection

To perform the CRISPR-based gene silencing, we developed Telo-KRAB: TeloHAEC stably expressing dCAS9-KRAB using a lenti-virus-based approach. The lentiviral vector lenti_dCas9-KRAB-MeCP2 was a gift from Andrea Califano (Addgene plasmid # 122205; http://n2t.net/addgene:122205; RRID: Addgene_122205). The viral vector was transduced to teloHAEC cells at MOI 20, screened by Blasticidin S HCL (Corning, Fisher Scientific# 15383671) selection (6 µg/ml) and stored in liquid nitrogen for future use. The guide RNAs were cloned into the pSPgRNA vector (pSPgRNA was a gift from Charles Gersbach, Addgene plasmid # 47108; http://n2t.net/addgene:47108; RRID: Addgene_47108) following standard protocol. A total of five gRNAs for region 2 (JCAD 2) and three gRNAs for JCAD 1 were cloned into the pSPgRNA vector and validated by restriction digestions and sanger sequencing. The Telo-KRAb cells were grown to 70-90% confluency before the transfection. The gRNA plasmids were mixed as pairs: one plasmid targeted the upstream, and other targeted the downstream region of the SNP of interest.

The control GFP plasmid (10% w/w) was also added to each combination to visualize the transfection efficiency microscopically. The cells were washed, trypsinized, and re-washed before the transfection reaction. A total of 50 000 cells were loaded along with plasmid mixture into each 10 µl tip of the Neon Transfection System kit. The optimized reaction condition used for Telo-KRAB cells was voltage: 1350V, width: 30ms, pulses: 1 pulse. Three electroporation reactions were performed for each treatment, and the cells were inoculated into a 12-well plate immediately after transfection. The cells were then grown for 24h to check the GFP expression as the sign of transfection efficiency. The samples for each reaction were collected after 48h of post-transfection. The RNA extraction was performed using Monarch® Total RNA Miniprep Kit. (NEB, cat T2010). To measure the extent of gene silencing, qPCR was performed for GAPDH and JCAD/KIAA1462 as described in 'CRISPR-RNP transfection'.

Prediction of TF binding motif disruption

To predict disruption of TF binding due to a SNP, the candidate SNPs were analyzed using all 94 high-confidence binding models that were published by Yan *et al.* (*70*) and derived by training deltaSVM models on *in vitro* protein–DNA binding data. The model is based on systematic measurements of the binding of 270 human transcription factors to 95,886 noncoding variants in the human genome using an ultra-high-throughput multiplex protein–DNA binding assay, SNP-SELEX. The resulting 828 million measurements of transcription factor–DNA interactions enabled the estimation of the relative affinity of these transcription factors to each variant *in vitro*. The original authors' recommended thresholds were used for determining sequence binding and allelic disruption. In addition, the consequence of the motif disruption was predicted based on the work of Kheradpour and Kellis (*71*) that is based on systematic motif analysis of 427 human ChIP-seq datasets.

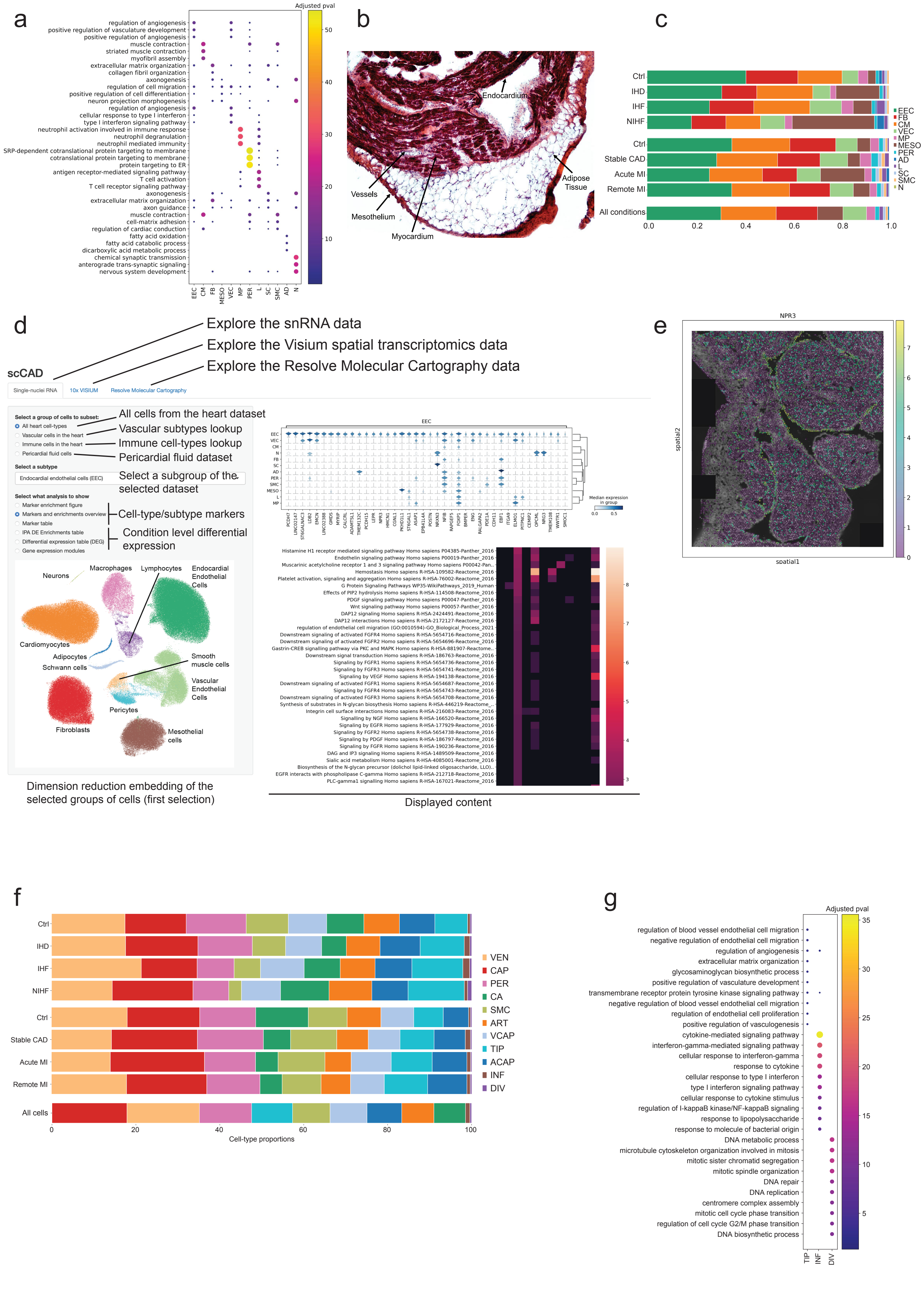

a
Adjusted pval
regulation of angiogenesis
positive regulation of vasculature development
positive regulation of angiogenesis
muscle contraction
striated muscle contraction
myofibril assembly
extracellular matrix organization
collagen fibril organization
axonogenesis
regulation of cell migration
positive regulation of cell differentiation
neuron projection morphogenesis
regulation of angiogenesis
cellular response to type I interferon
type I interferon signaling pathway
neutrophil activation involved in immune response
neutrophil degranulation
neutrophil mediated immunity
SRP-dependent cotranslational protein targeting to membrane
cotranslational protein targeting to membrane
protein targeting to ER
antigen receptor-mediated signaling pathway
T cell activation
T cell receptor signaling pathway
axonogenesis
extracellular matrix organization
axon guidance
muscle contraction
cell-matrix adhesion
regulation of cardiac conduction
fatty acid oxidation
fatty acid catabolic process
dicarboxylic acid metabolic process
chemical synaptic transmission
anterograde trans-synaptic signaling
nervous system development
EEC CM FB MESO VEC MP PER L SC SMC AD N
b
Endocardium
Vessels
Mesothelium
Myocardium
Adipose Tissue
c
Ctrl
IHD
IHF
NIHF
Ctrl
Stable CAD
Acute MI
Remote MI
All conditions
EEC FB CM VEC MP MESO PER AD L SC SMC N
d
Explore the snRNA data
Explore the Visium spatial transcriptomics data
Explore the Resolve Molecular Cartography data
scCAD
Single-nuclei RNA
10x VISIUM
Resolve Molecular Cartography
Select a group of cells to subset:
All heart cell-types
Vascular cells in the heart
Immune cells in the heart
Pericardial fluid cells
All cells from the heart dataset
Vascular subtypes lookup
Immune cell-types lookup
Pericardial fluid dataset
Select a subtype
Endocardial endothelial cells (EEC)
Select a subgroup of the selected dataset
Select what analysis to show
Marker enrichment figure
Markers and enrichments overview
Marker table
IPA DE Enrichments table
Differential expression table (DEG)
Gene expression modules
Cell-type/subtype markers
Condition level differential expression
Neurons
Macrophages
Lymphocytes
Endocardial Endothelial Cells
Cardiomyocytes
Adipocytes
Schwann cells
Smooth muscle cells
Vascular Endothelial Cells
Pericytes
Fibroblasts
Mesothelial cells
Dimension reduction embedding of the selected groups of cells (first selection)
EEC
Median expression in group
Displayed content
e
NPR3
spatial2
spatial1
f
Ctrl
IHD
IHF
NIHF
Ctrl
Stable CAD
Acute MI
Remote MI
All cells
VEN CAP PER CA SMC ART VCAP TIP ACAP INF DIV
Cell-type proportions
g
Adjusted pval
regulation of blood vessel endothelial cell migration
negative regulation of endothelial cell migration
regulation of angiogenesis
extracellular matrix organization
glycosaminoglycan biosynthetic process
positive regulation of vasculature development
transmembrane receptor protein tyrosine kinase signaling pathway
negative regulation of blood vessel endothelial cell migration
regulation of endothelial cell proliferation
positive regulation of vasculogenesis
cytokine-mediated signaling pathway
interferon-gamma-mediated signaling pathway
cellular response to interferon-gamma
response to cytokine
cellular response to type I interferon
type I interferon signaling pathway
cellular response to cytokine stimulus
regulation of I-kappaB kinase/NF-kappaB signaling
response to lipopolysaccharide
response to molecule of bacterial origin
DNA metabolic process
microtubule cytoskeleton organization involved in mitosis
mitotic sister chromatid segregation
mitotic spindle organization
DNA repair
DNA replication
centromere complex assembly
mitotic cell cycle phase transition
regulation of cell cycle G2/M phase transition
DNA biosynthetic process
TIP INF DIV

**Fig. S1.**

**a.** Top 3 enriched gene sets (GO Biological Process) for top 1000 genes with FDR $< 0.05$ for each cell type. Dot color and size is -log10(adjusted p-value). Only significant (adjusted p-value $< 0.05$) enrichments are shown. **b.** Hematoxylin-eosin-stained histology slide of a right atrial sample included in the dataset. **c.** Proportion of the main cell types, per condition. **d.** Interactive website overview. An interactive website is accessible at **website**. It provides access to the analyzed snRNA-seq and spatial transcriptomics data (both Visium and Resolve Molecular Cartography) presented in this study. The user can select the ensemble of cells to focus on (all heart cells, vascular heart cells, immune heart cells, pericardial fluid data) and explore different information types (1. Gene set enrichment of subtype/cell-type marker genes 2. Subtype/cell-type marker gene overview. 3. Subtype/cell-type marker genes table. 4. IPA enrichment table from differential expression result of a selected condition in the subtype/cell-type. 5. Differential expression result table of a selected condition in the subtype/cell-type. 6. Gene expression module analysis in the selected subtype/cell-type). **e.** Spatial expression of *NPR3* on a Resolve molecular cartography section of the right atrium. **f.** Proportion of vascular cells, per condition. **g**. Top 10 enriched gene sets (GO Biological Process) for top 200 genes with FDR $< 0.05$ in each subtype, for tip cells (TIP), inflammatory endothelial cells (INF) and dividing endothelial cells (DIV). Dot color and size is -log10(adjusted enrichment p-value). Only significant (adjusted p-value $< 0.05$) enrichments are shown.

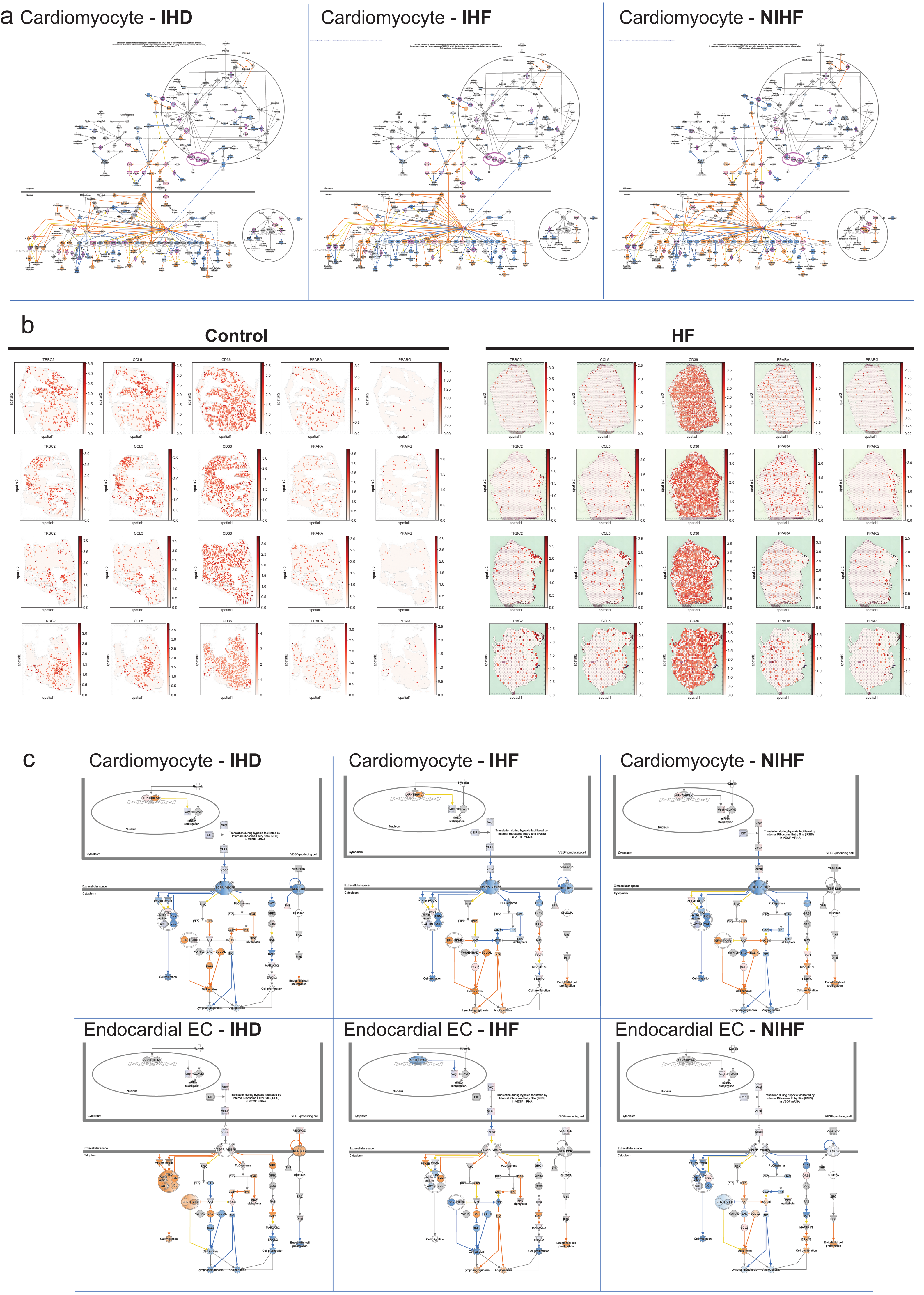
a
Cardiomyocyte - IHD
Cardiomyocyte - IHF
Cardiomyocyte - NIHF
b
Control
HF
TRBC2
CCL5
CD36
PPARA
PPARG
spatial1
spatial2
c
Cardiomyocyte - IHD
Cardiomyocyte - IHF
Cardiomyocyte - NIHF
Endocardial EC - IHD
Endocardial EC - IHF
Endocardial EC - NIHF
Hypoxia
Nucleus
Cytoplasm
VEGF-producing cell
Translation during hypoxia facilitated by Internal Ribosome Entry Site (IRES) in VEGF mRNA
Extracellular space
Cell migration
Cell survival
Cell proliferation
Endothelial cell proliferation
Lymphangiogenesis
Angiogenesis

**Fig. S2.**

**a.** Sirtuin signaling pathway from IPA in cardiomyocytes in IHD, IHF, and NIHF. Colors indicate measured and inferred expression/activity changes. Blue stands for downregulation or inhibition, orange/red for upregulation. Yellow line indicates inconsistent changes and purple highlights measured significant changes in the dataset. **b.** Expression of *TRBC2*, *CCL5*, *CD36*, *PPARA*, *PPARG* on Visium slides, from a patient with heart failure and a control sample ($n$=4). **c.** VEGF signaling pathway from IPA in cardiomyocytes and endocardial endothelial cells in IHD, IHF, and NIHF. Blue stands for downregulation or inhibition of the molecule, orange/red for upregulation. Yellow line indicates inconsistent changes.

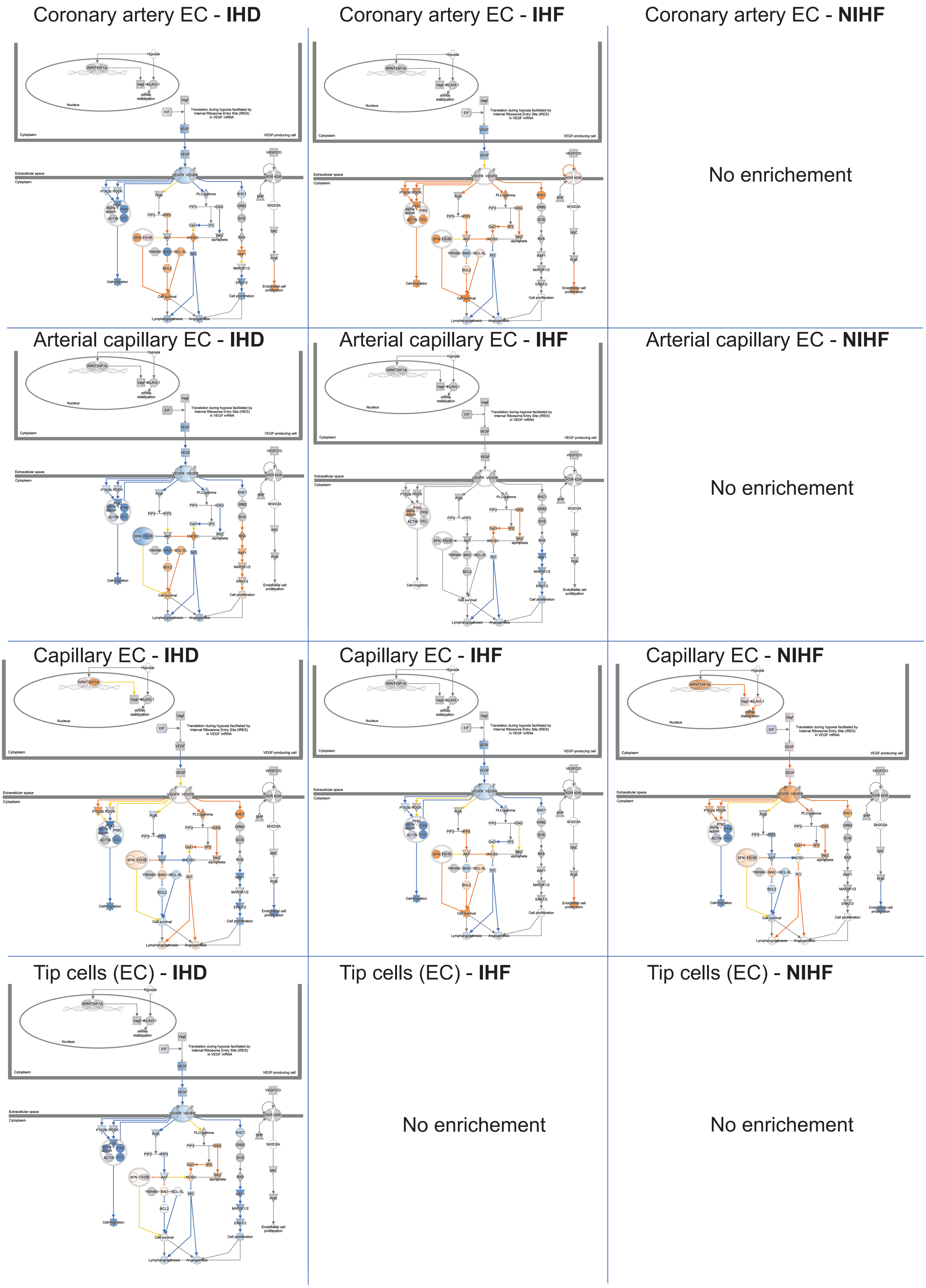

Coronary artery EC - **IHD**
Coronary artery EC - **IHF**
Coronary artery EC - **NIHF**
No enrichement
Arterial capillary EC - **IHD**
Arterial capillary EC - **IHF**
Arterial capillary EC - **NIHF**
No enrichement
Capillary EC - **IHD**
Capillary EC - **IHF**
Capillary EC - **NIHF**
Tip cells (EC) - **IHD**
Tip cells (EC) - **IHF**
Tip cells (EC) - **NIHF**
No enrichement
No enrichement

**Fig. S3.**

VEGF signaling pathway from IPA in endothelial subtypes. Blue stands for downregulation or inhibition, orange/red for upregulation. Yellow line indicates inconsistent changes.

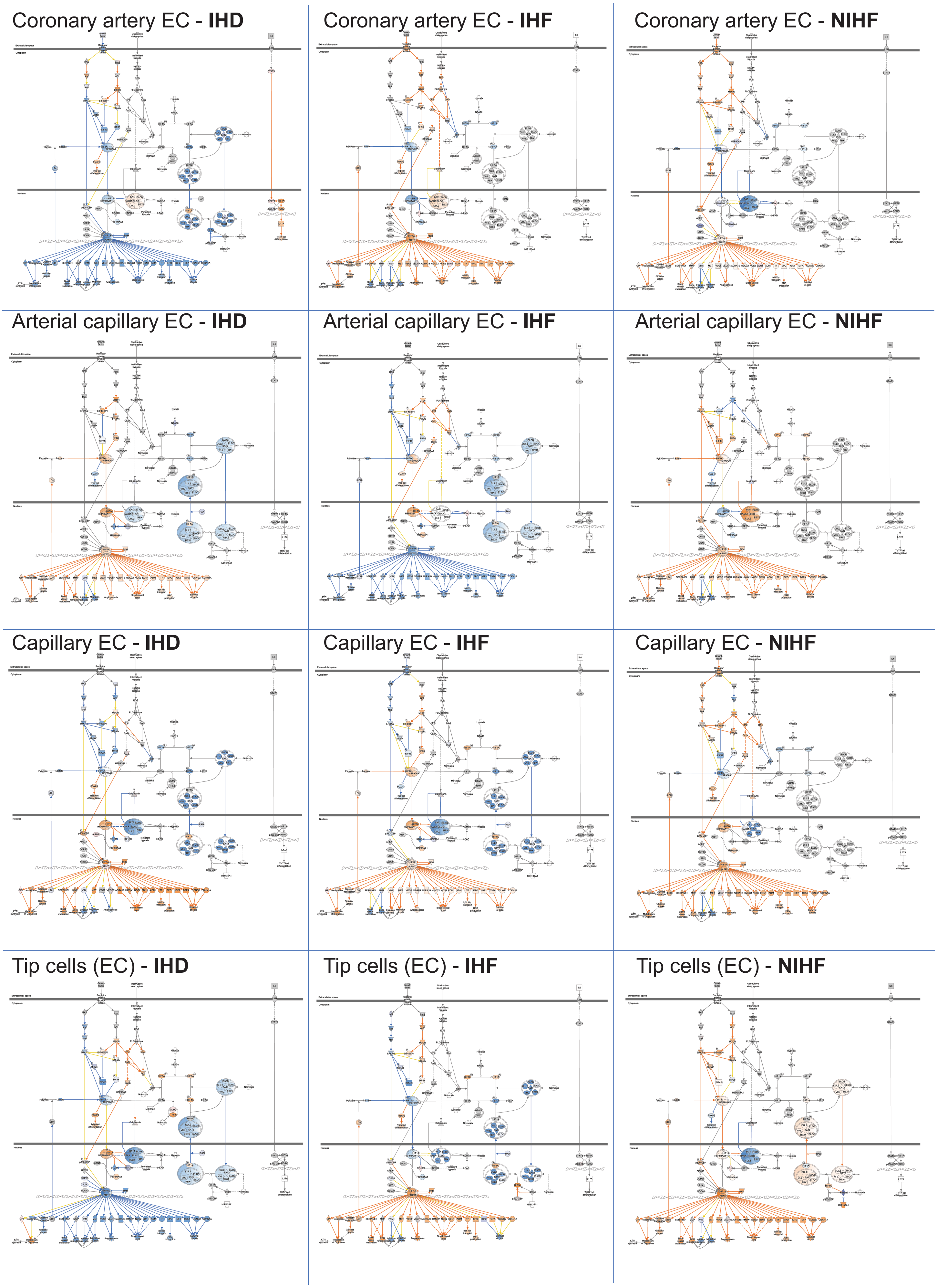
Coronary artery EC - **IHD**
Coronary artery EC - **IHF**
Coronary artery EC - **NIHF**
Arterial capillary EC - **IHD**
Arterial capillary EC - **IHF**
Arterial capillary EC - **NIHF**
Capillary EC - **IHD**
Capillary EC - **IHF**
Capillary EC - **NIHF**
Tip cells (EC) - **IHD**
Tip cells (EC) - **IHF**
Tip cells (EC) - **NIHF**

**Fig. S4.**

HIF signaling pathway from IPA in endothelial subtypes in IHD, IHF, and NIHF. Blue stands for downregulation or inhibition, orange/red for upregulation. Yellow line indicates inconsistent changes.

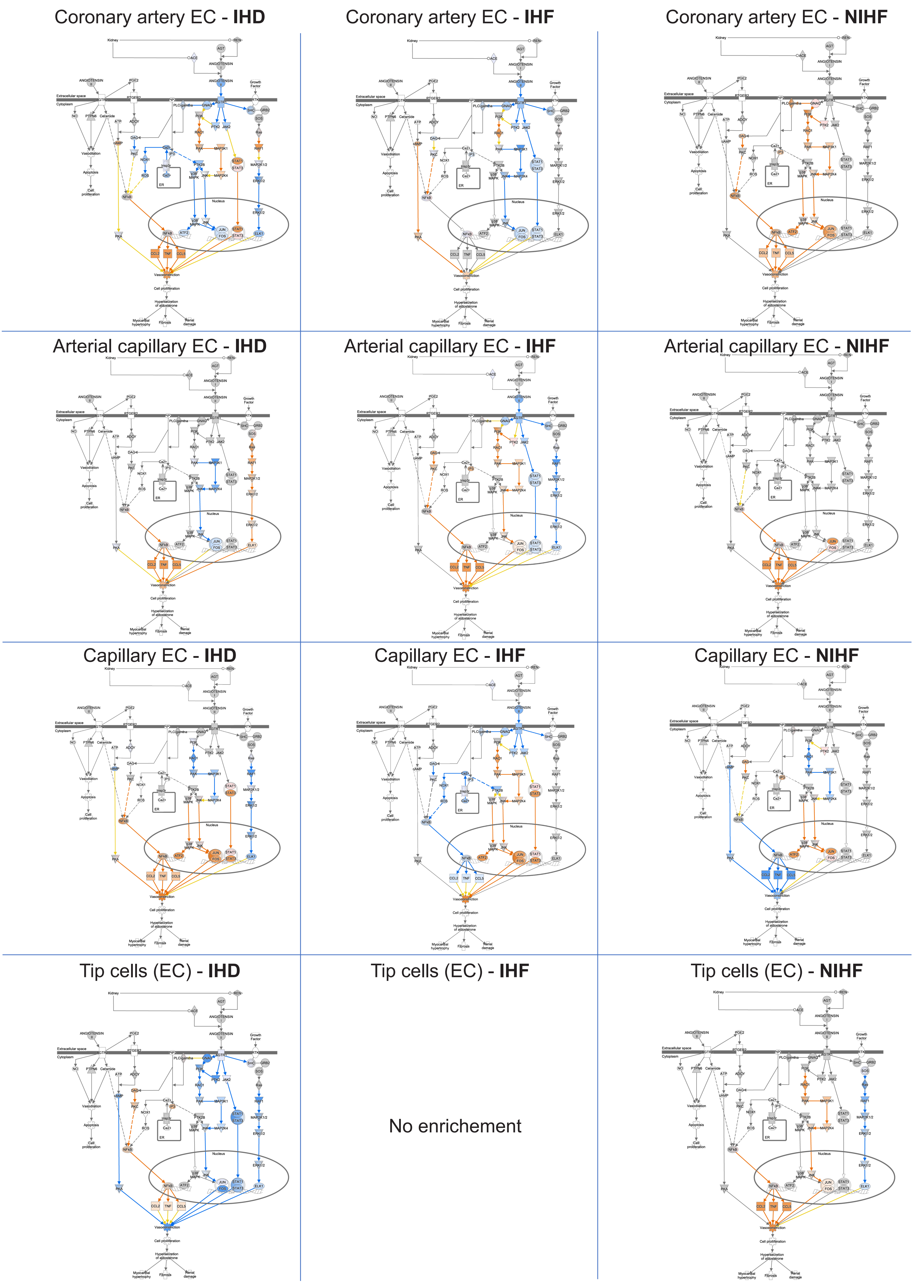
Coronary artery EC - IHD
Coronary artery EC - IHF
Coronary artery EC - NIHF
Arterial capillary EC - IHD
Arterial capillary EC - IHF
Arterial capillary EC - NIHF
Capillary EC - IHD
Capillary EC - IHF
Capillary EC - NIHF
Tip cells (EC) - IHD
Tip cells (EC) - IHF
No enrichement
Tip cells (EC) - NIHF

**Fig. S5.**

Renin signaling pathway from IPA in endothelial subtypes in IHD, IHF, and NIHF. Blue stands for downregulation or inhibition, orange/red for upregulation. Yellow line indicates inconsistent changes.

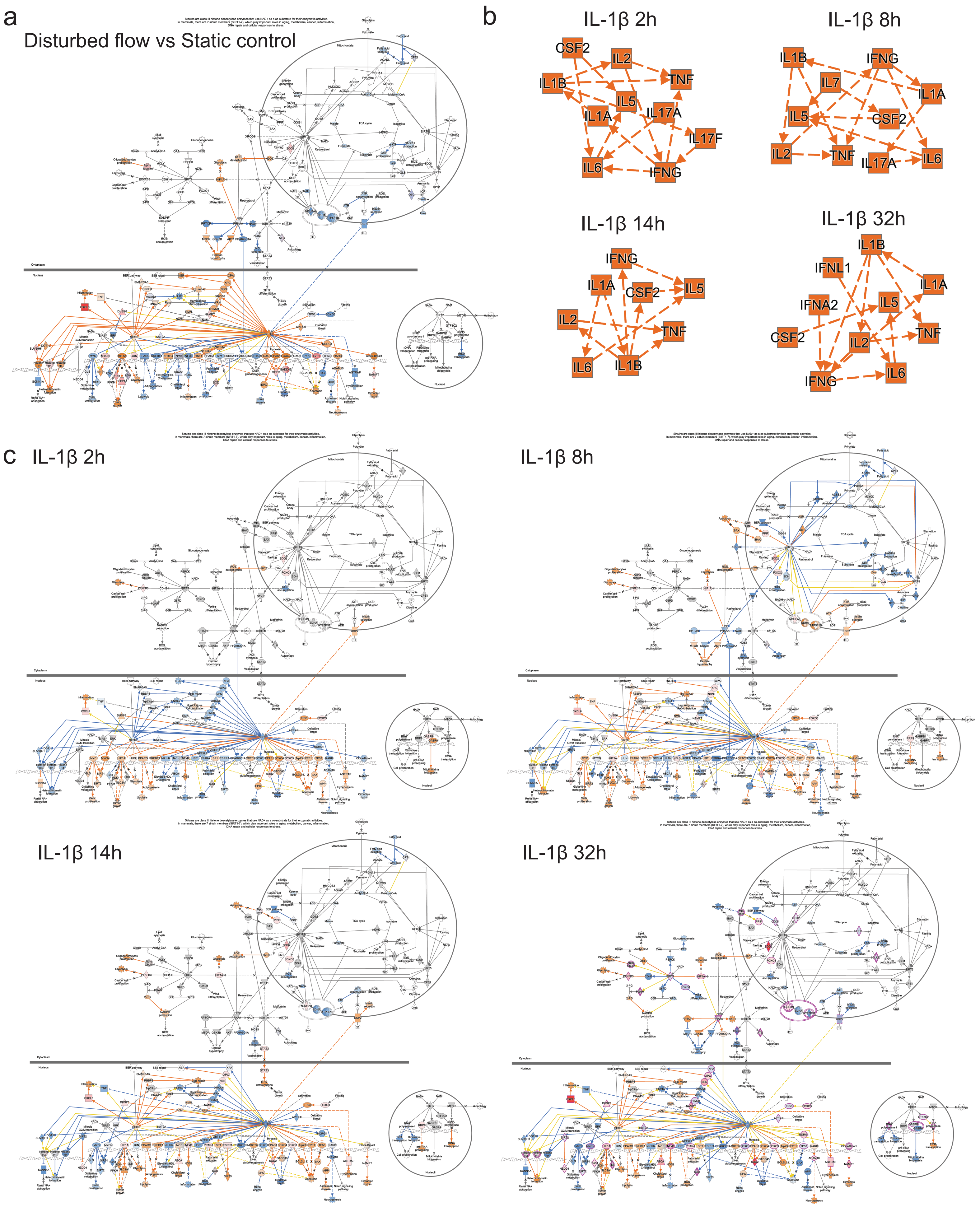

a
Disturbed flow vs Static control
b
IL-1β 2h
CSF2
IL2
TNF
IL1B
IL5
IL1A
IL17A
IL17F
IL6
IFNG
IL-1β 8h
IL1B
IFNG
IL7
IL1A
IL5
CSF2
IL2
TNF
IL17A
IL6
IL-1β 14h
IFNG
IL1A
CSF2
IL5
IL2
TNF
IL6
IL1B
IL-1β 32h
IL1B
IFNL1
IL1A
IFNA2
IL5
CSF2
TNF
IL2
IFNG
IL6
c
IL-1β 2h
IL-1β 8h
IL-1β 14h
IL-1β 32h

**Fig. S6.**

**a.** Sirtuin signaling pathway from IPA in HUVEC cells under atheroprone (disturbed) flow versus static cells (*N*=3). Blue stands for downregulation or inhibition, orange/red for upregulation. Yellow line indicates inconsistent changes. **b.** IL-1β network from IPA in stimulated HAEC cells. The cells were treated for 2, 8, 14, or 32h with IL-1β. The resulting RNA-seq data were analyzed with IPA (Qiagen) to construct the upstream regulator networks from the graphical summaries. **c.** Sirtuin signaling pathway from IPA in IL-1β-treated HAECs after 2, 8, 14, and 32h of treatment. Blue stands for downregulation or inhibition, orange/red for upregulation. Yellow line indicates inconsistent changes and purple highlights measured significant changes in the dataset. All signaling pathways were constructed with IPA (Qiagen) from either core or comparison analysis.

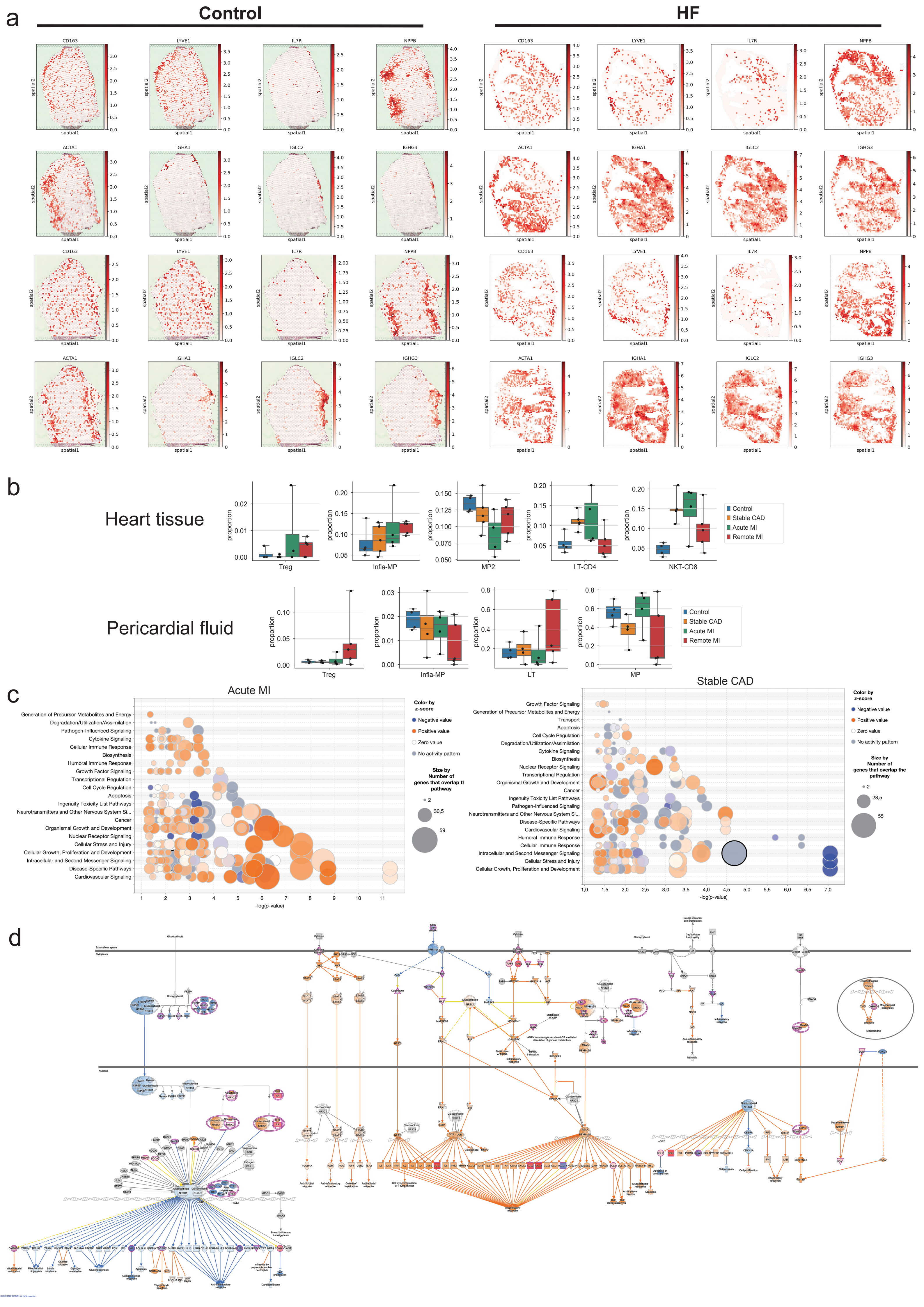
a
Control
HF
CD163
LYVE1
IL7R
NPPB
ACTA1
IGHA1
IGLC2
IGHG3
spatial1
spatial2
b
Heart tissue
Pericardial fluid
proportion
Treg
Infla-MP
MP2
LT-CD4
NKT-CD8
LT
MP
Control
Stable CAD
Acute MI
Remote MI
c
Acute MI
Stable CAD
Generation of Precursor Metabolites and Energy
Degradation/Utilization/Assimilation
Pathogen-Influenced Signaling
Cytokine Signaling
Cellular Immune Response
Biosynthesis
Humoral Immune Response
Growth Factor Signaling
Transcriptional Regulation
Cell Cycle Regulation
Apoptosis
Ingenuity Toxicity List Pathways
Neurotransmitters and Other Nervous System Si...
Cancer
Organismal Growth and Development
Nuclear Receptor Signaling
Cellular Stress and Injury
Cellular Growth, Proliferation and Development
Intracellular and Second Messenger Signaling
Disease-Specific Pathways
Cardiovascular Signaling
Transport
-log(p-value)
Color by z-score
Negative value
Positive value
Zero value
No activity pattern
Size by Number of genes that overlap the pathway
d
Extracellular space
Cytoplasm
Nucleus
Inflammatory response

**Fig. S7.**

**a.** Expression of *CD163*, *LYVE1*, *IL7R* on Visium slides, from a patient with heart failure and a control sample (*n*=4). **b.** Proportion of selected immune cells across conditions in heart tissue and pericardial fluid. Whiskers show the maximum and minimum values, except for outliers (more than 1.5 times the interquartile). **c.** Enrichment of general terms and pathway sharing between terms in macrophages (MP) in stable CAD and acute MI from IPA. A pathway shared between the terms is shown as a circle of the same size and color on the same vertical line between the terms. **d.** Glucocorticoid signaling pathway from IPA in MPs in stable CAD compared to control. Blue stands for downregulation or inhibition, orange/red for upregulation. Yellow line indicates inconsistent changes and purple highlights measured significant changes in the dataset.

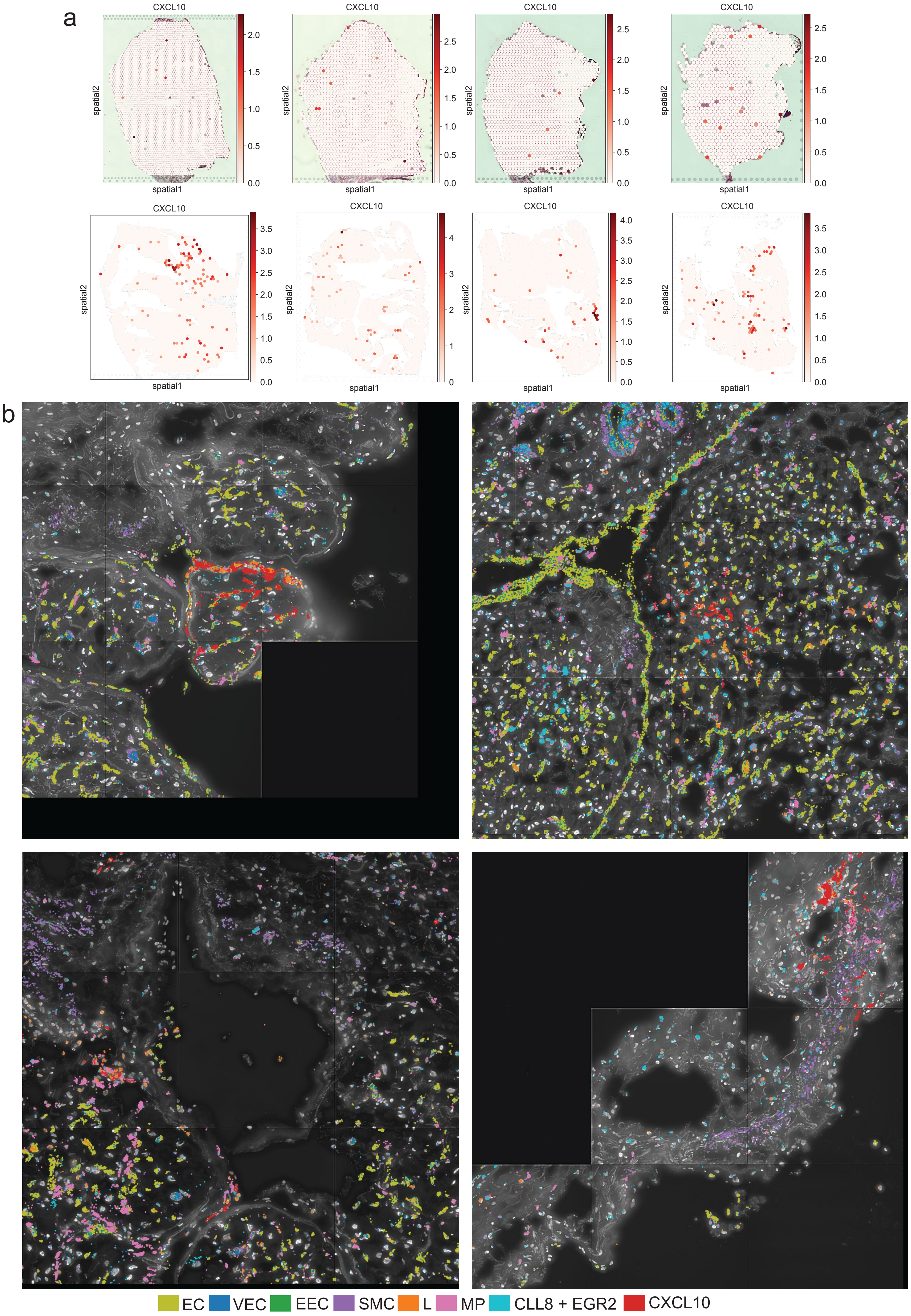
a
CXCL10
spatial1
spatial2
b
EC
VEC
EEC
SMC
L
MP
CLL8 + EGR2
CXCL10

**Fig. S8.**

**a.** Expression of *CXCL10* on Visium slides, from a patient with heart failure and a control sample (*n*=4). **b.** Spatial images (Resolve Molecular Cartography) for EC (*EMCN, ERG, PECAM1, CDH5, VWF*), VEC (*DKK2, ENPP2, PCSK5, CYYR1*), EEC (*PCDH7*), SMC (*NTRK3, MRVII*), L (*BCL11B, CD247, SKAP1, THEMIS*), MP (*CD163, MRC1, F13A1, MS4A6A*), inflammatory (*CCL8* and *EGR2*), and *CXCL10* gene expression.

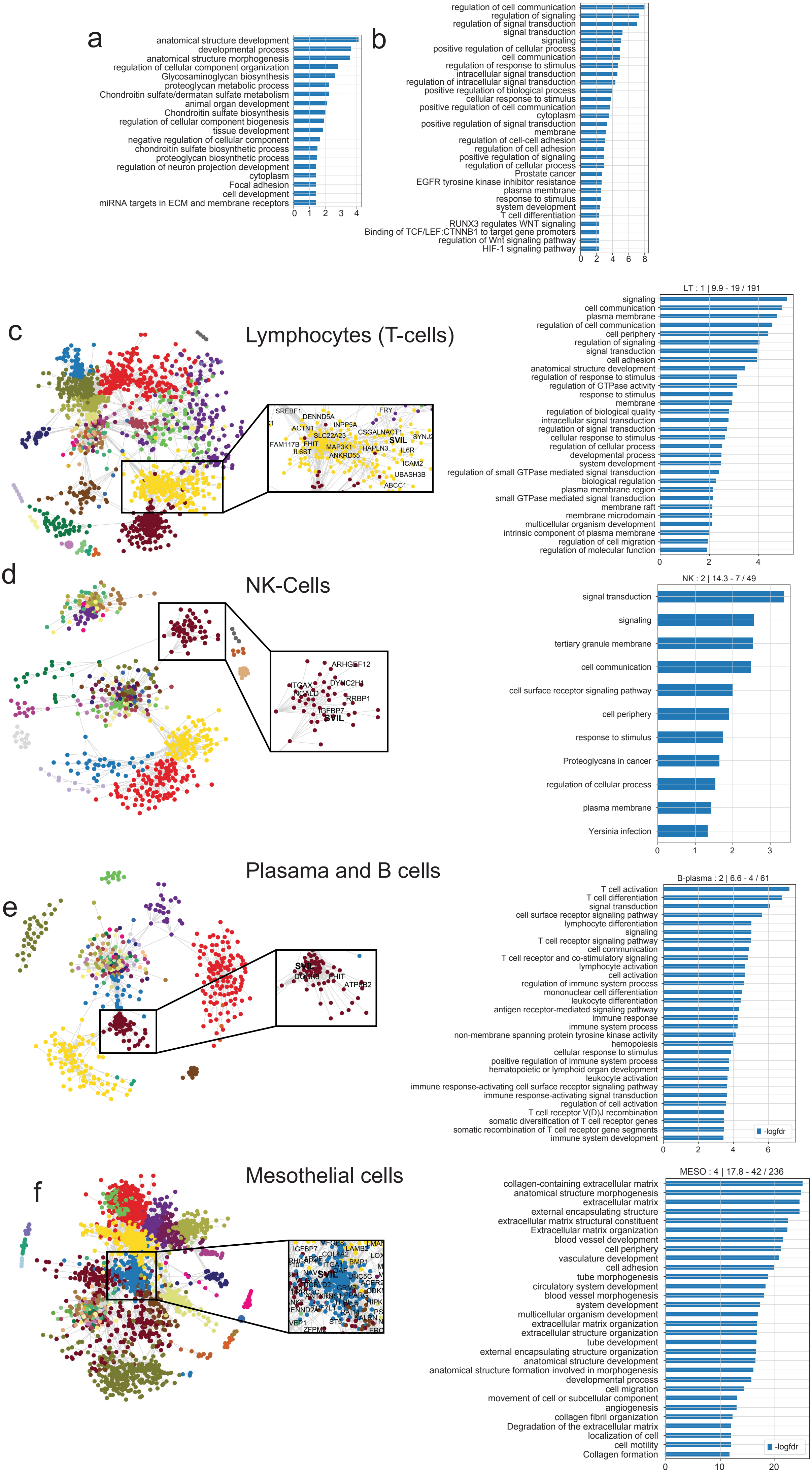
a
anatomical structure development
developmental process
anatomical structure morphogenesis
regulation of cellular component organization
Glycosaminoglycan biosynthesis
proteoglycan metabolic process
Chondroitin sulfate/dermatan sulfate metabolism
animal organ development
Chondroitin sulfate biosynthesis
regulation of cellular component biogenesis
tissue development
negative regulation of cellular component
chondroitin sulfate biosynthetic process
proteoglycan biosynthetic process
regulation of neuron projection development
cytoplasm
Focal adhesion
cell development
miRNA targets in ECM and membrane receptors
0 1 2 3 4
b
regulation of cell communication
regulation of signaling
regulation of signal transduction
signal transduction
signaling
positive regulation of cellular process
cell communication
regulation of response to stimulus
intracellular signal transduction
regulation of intracellular signal transduction
positive regulation of biological process
cellular response to stimulus
positive regulation of cell communication
cytoplasm
positive regulation of signal transduction
membrane
regulation of cell-cell adhesion
regulation of cell adhesion
positive regulation of signaling
regulation of cellular process
Prostate cancer
EGFR tyrosine kinase inhibitor resistance
plasma membrane
response to stimulus
system development
T cell differentiation
RUNX3 regulates WNT signaling
Binding of TCF/LEF:CTNNB1 to target gene promoters
regulation of Wnt signaling pathway
HIF-1 signaling pathway
0 2 4 6 8
c
Lymphocytes (T-cells)
SREBF1
DENND5A
FRY
INPP5A
ACTN1
CSGALNACT1
SLC22A23
SYNJ2
FAM117B
FHIT
SVIL
MAP3K1
HAPLN3
IL6ST
ANKRD55
IL6R
ICAM2
UBASH3B
ABCC1
LT : 1 | 9.9 - 19 / 191
signaling
cell communication
plasma membrane
regulation of cell communication
cell periphery
regulation of signaling
signal transduction
cell adhesion
anatomical structure development
regulation of response to stimulus
regulation of GTPase activity
response to stimulus
membrane
regulation of biological quality
intracellular signal transduction
regulation of signal transduction
cellular response to stimulus
regulation of cellular process
developmental process
system development
regulation of small GTPase mediated signal transduction
biological regulation
plasma membrane region
small GTPase mediated signal transduction
membrane raft
membrane microdomain
multicellular organism development
intrinsic component of plasma membrane
regulation of cell migration
regulation of molecular function
0 2 4
d
NK-Cells
ARHGEF12
ITGAX
DYNC2H1
HCALD
RRBP1
IGFBP7
SVIL
NK : 2 | 14.3 - 7 / 49
signal transduction
signaling
tertiary granule membrane
cell communication
cell surface receptor signaling pathway
cell periphery
response to stimulus
Proteoglycans in cancer
regulation of cellular process
plasma membrane
Yersinia infection
0 1 2 3
Plasama and B cells
e
SVIL
DOCK9
FHIT
ATP8B2
B-plasma : 2 | 6.6 - 4 / 61
T cell activation
T cell differentiation
signal transduction
cell surface receptor signaling pathway
lymphocyte differentiation
signaling
T cell receptor signaling pathway
cell communication
T cell receptor and co-stimulatory signaling
lymphocyte activation
cell activation
regulation of immune system process
mononuclear cell differentiation
leukocyte differentiation
antigen receptor-mediated signaling pathway
immune response
immune system process
non-membrane spanning protein tyrosine kinase activity
hemopoiesis
cellular response to stimulus
positive regulation of immune system process
hematopoietic or lymphoid organ development
leukocyte activation
immune response-activating cell surface receptor signaling pathway
immune response-activating signal transduction
regulation of cell activation
T cell receptor V(D)J recombination
somatic diversification of T cell receptor genes
somatic recombination of T cell receptor gene segments
immune system development
-logfdr
0 2 4 6
Mesothelial cells
f
IGFBP7
LAMB2
COL4A2
BMP1
SVIL
ITGA1
ZFPM
MESO : 4 | 17.8 - 42 / 236
collagen-containing extracellular matrix
anatomical structure morphogenesis
extracellular matrix
external encapsulating structure
extracellular matrix structural constituent
Extracellular matrix organization
blood vessel development
cell periphery
vasculature development
cell adhesion
tube morphogenesis
circulatory system development
blood vessel morphogenesis
system development
multicellular organism development
extracellular matrix organization
extracellular structure organization
tube development
external encapsulating structure organization
anatomical structure development
anatomical structure formation involved in morphogenesis
developmental process
cell migration
movement of cell or subcellular component
angiogenesis
collagen fibril organization
Degradation of the extracellular matrix
localization of cell
cell motility
Collagen formation
-logfdr
0 10 20

**Fig. S9.**

**a.** Gene set enrichment with Enrichr for the gene-gene co-expression module containing *SVIL* in SMC. **b.** Gene set enrichment with Enrichr of the gene-gene co-expression module containing *SVIL* in pericardial fluid cells. **c.** Gene-gene co-expression modules within Pericardial fluid T lymphocytes, NK cells, B and plasma cells and Mesothelial cells, in addition to a zoom in view of *SVIL*-containing modules, with their associated gene set enrichments from Enrichr.

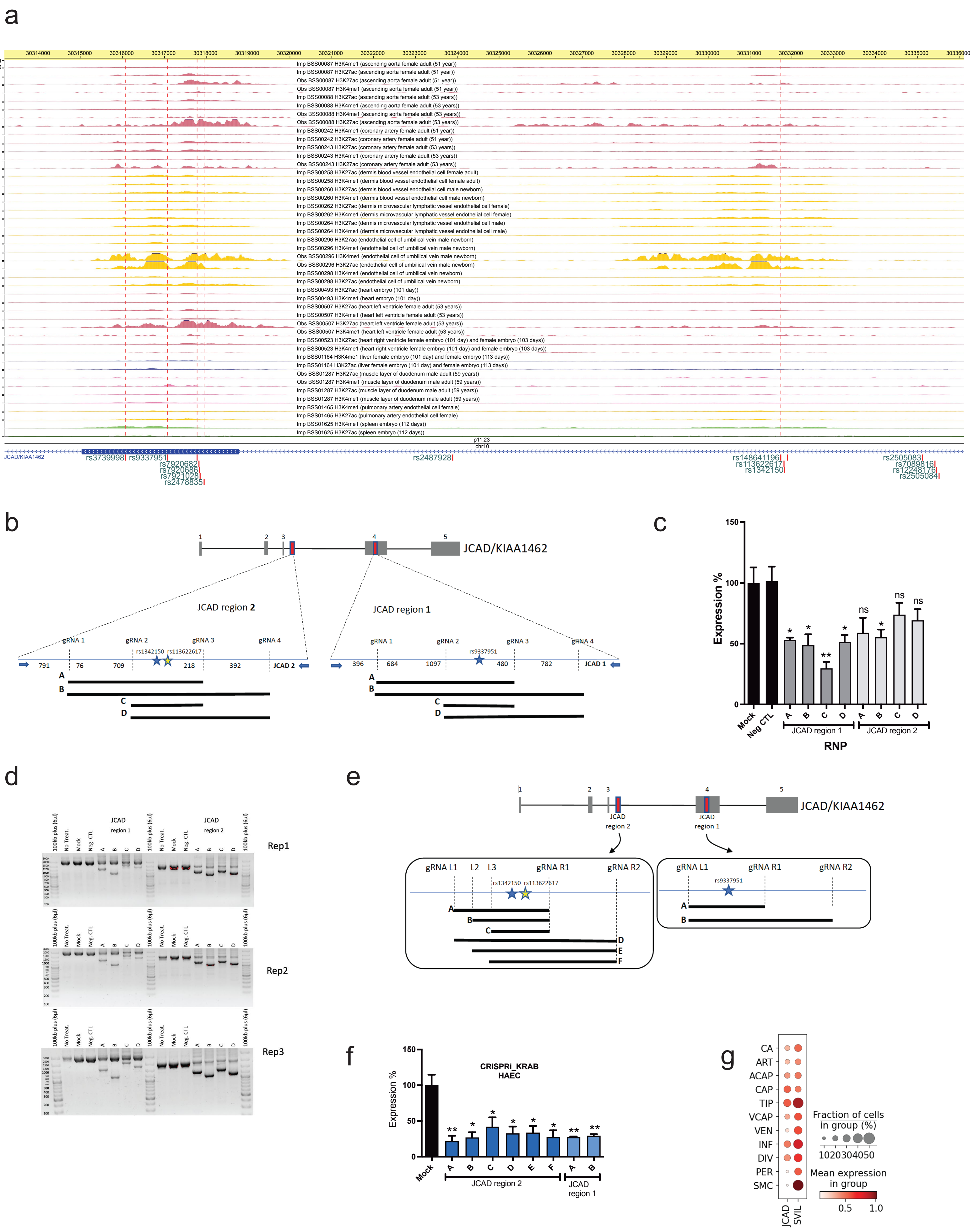

a
Imp BSS00087 H3K4me1 (ascending aorta female adult (51 year))
Imp BSS00087 H3K27ac (ascending aorta female adult (51 year))
Obs BSS00087 H3K27ac (ascending aorta female adult (51 year))
Obs BSS00087 H3K4me1 (ascending aorta female adult (51 year))
Imp BSS00088 H3K27ac (ascending aorta female adult (53 years))
Imp BSS00088 H3K4me1 (ascending aorta female adult (53 years))
Obs BSS00088 H3K4me1 (ascending aorta female adult (53 years))
Obs BSS00088 H3K27ac (ascending aorta female adult (53 years))
Imp BSS00242 H3K4me1 (coronary artery female adult (51 year))
Imp BSS00242 H3K27ac (coronary artery female adult (51 year))
Imp BSS00243 H3K27ac (coronary artery female adult (53 years))
Imp BSS00243 H3K4me1 (coronary artery female adult (53 years))
Obs BSS00243 H3K27ac (coronary artery female adult (53 years))
Imp BSS00258 H3K27ac (dermis blood vessel endothelial cell female adult)
Imp BSS00258 H3K4me1 (dermis blood vessel endothelial cell female adult)
Imp BSS00260 H3K27ac (dermis blood vessel endothelial cell male newborn)
Imp BSS00260 H3K4me1 (dermis blood vessel endothelial cell male newborn)
Imp BSS00262 H3K27ac (dermis microvascular lymphatic vessel endothelial cell female)
Imp BSS00262 H3K4me1 (dermis microvascular lymphatic vessel endothelial cell female)
Imp BSS00264 H3K27ac (dermis microvascular lymphatic vessel endothelial cell male)
Imp BSS00264 H3K4me1 (dermis microvascular lymphatic vessel endothelial cell male)
Imp BSS00296 H3K27ac (endothelial cell of umbilical vein male newborn)
Imp BSS00296 H3K4me1 (endothelial cell of umbilical vein male newborn)
Obs BSS00296 H3K4me1 (endothelial cell of umbilical vein male newborn)
Obs BSS00296 H3K27ac (endothelial cell of umbilical vein male newborn)
Imp BSS00298 H3K4me1 (endothelial cell of umbilical vein newborn)
Imp BSS00298 H3K27ac (endothelial cell of umbilical vein newborn)
Imp BSS00493 H3K27ac (heart embryo (101 day))
Imp BSS00493 H3K4me1 (heart embryo (101 day))
Imp BSS00507 H3K27ac (heart left ventricle female adult (53 years))
Imp BSS00507 H3K4me1 (heart left ventricle female adult (53 years))
Obs BSS00507 H3K27ac (heart left ventricle female adult (53 years))
Obs BSS00507 H3K4me1 (heart left ventricle female adult (53 years))
Imp BSS00523 H3K27ac (heart right ventricle female embryo (101 day) and female embryo (103 days))
Imp BSS00523 H3K4me1 (heart right ventricle female embryo (101 day) and female embryo (103 days))
Imp BSS01164 H3K4me1 (liver female embryo (101 day) and female embryo (113 days))
Imp BSS01164 H3K27ac (liver female embryo (101 day) and female embryo (113 days))
Obs BSS01287 H3K27ac (muscle layer of duodenum male adult (59 years))
Obs BSS01287 H3K4me1 (muscle layer of duodenum male adult (59 years))
Imp BSS01287 H3K27ac (muscle layer of duodenum male adult (59 years))
Imp BSS01287 H3K4me1 (muscle layer of duodenum male adult (59 years))
Imp BSS01465 H3K4me1 (pulmonary artery endothelial cell female)
Imp BSS01465 H3K27ac (pulmonary artery endothelial cell female)
Imp BSS01625 H3K4me1 (spleen embryo (112 days))
Imp BSS01625 H3K27ac (spleen embryo (112 days))
p11.23
chr10
JCAD/KIAA1462
rs3739998 rs9337951
rs7920682 rs7920686 rs7921028 rs2478835
rs2487928
rs148641196 rs113622617 rs1342150
rs2505083 rs7089816 rs12248176 rs2505084
b
JCAD/KIAA1462
JCAD region 2
JCAD region 1
gRNA 1 gRNA 2 gRNA 3 gRNA 4
rs1342150 rs113622617
rs9337951
791 76 709 218 392 JCAD 2
396 684 1097 480 782 JCAD 1
A B C D
c
Expression %
Mock Neg CTL
JCAD region 1
JCAD region 2
RNP
ns
d
100kb plus (6µl)
No Treat.
Mock
Neg. CTL
JCAD region 1
JCAD region 2
Rep1
Rep2
Rep3
e
JCAD/KIAA1462
JCAD region 2
JCAD region 1
gRNA L1 L2 L3 gRNA R1 gRNA R2
gRNA L1 gRNA R1 gRNA R2
rs1342150 rs113622617
rs9337951
f
CRISPRi_KRAB
HAEC
Expression %
Mock
JCAD region 2
JCAD region 1
g
CA ART ACAP CAP TIP VCAP VEN INF DIV PER SMC
Fraction of cells in group (%)
1020304050
Mean expression in group
0.5 1.0
JCAD SVIL

**Fig. S10.**

**a.** Browser shot of the rs9337951 locus on chromosome 10 with histone marker signals for aorta, endothelial cells, heart, liver, muscle, and spleen. **b**. Schematic describing the regions targeted for CRISPR-mediated deletion using different combinations of two guide RNAs (gRNAs within the *JCAD*/*SVIL* locus. **c.** The effect of CRISPR deletion on *JCAD* gene expression. **d.** Confirmation of the success and efficiency of deletion using PCR. **e.** Schematic describing the regions targeted for CRISPR-mediated inhibition using different combinations of two gRNAs **f.** The effect of CRISPR inhibition on *JCAD* gene expression. **g.** *JCAD* and *SVIL* expression in tissue cells by snRNA-seq.

**Table S1.**

Results for allelic activity reporter assay

| | | teloHAEC 0h | | teloHAEC 6h Il1b | | teloHAEC 24h Il1b | | HASMC chol | | |
|---|---|---|---|---|---|---|---|---|---|---|
| RS Number | Position (GRCh37) | logFC | FDR | logFC | FDR | logFC | FDR | logFC | FDR | variant type |
| rs2478839 | chr10:30306804 | 0,029497 | 0,89317 | -0,22841 | 0,594769 | 0,068412 | 0,895356 | 0,326823 | 0,203501 | common |
| rs12762440 | chr10:30306879 | 0,503904 | 0,12795 | 0,584236 | 0,479108 | -0,21375 | 0,566648 | -0,50645 | 0,101235 | rare |
| rs9337951 | chr10:30317073 | -0,70934 | 0,0661 | -0,22455 | 0,555735 | -0,08465 | 0,895356 | 0,471101 | 0,201433 | common |
| rs531337994 | chr10:30317781 | 0,721114 | 0,0661 | 0,307885 | 0,488862 | 0,871545 | 0,16561 | 1,106221 | 0,004756 | rare |
| rs7920682 | chr10:30317826 | -0,44092 | 0,17706 | 0,519276 | 0,479108 | -0,27119 | 0,56637 | -0,58955 | 0,115174 | common |
| rs7920686 | chr10:30317838 | -0,44092 | 0,17706 | 0,519276 | 0,479108 | -0,27119 | 0,56637 | -0,58955 | 0,115174 | common |
| rs7921028 | chr10:30317853 | -0,50678 | 0,33249 | 1,147921 | 0,479108 | -0,40207 | 0,56637 | -0,1748 | 0,75763 | rare |
| rs2478835 | chr10:30317949 | 0,256671 | 0,36864 | -0,24893 | 0,549716 | 0,61482 | 0,264878 | 0,504799 | 0,101235 | common |
| rs2487928 | chr10:30323892 | -0,16943 | 0,42384 | 0,312584 | 0,479108 | -0,02049 | 0,912102 | -0,26084 | 0,2621 | common |
| rs148641196 | chr10:30331736 | -0,92915 | 0,00225 | -1,3914 | 0,001085 | 0,108742 | 0,895356 | -1,05572 | 0,004181 | rare |
| rs113622617 | chr10:30331813 | 0,383709 | 0,12795 | 0,696887 | 0,042102 | 0,774711 | 0,49638 | 0,681045 | 0,091074 | rare |
| rs1342150 | chr10:30331829 | -0,4658 | 0,0661 | -1,0913 | 0,003272 | -0,68255 | 0,49638 | -0,83159 | 0,004756 | common |
| rs193042870 | chr10:30331895 | 0,353719 | 0,18304 | 0,954916 | 0,041686 | 1,121558 | 0,332584 | -0,4443 | 0,116393 | rare |
| rs7089816 | chr10:30335422 | -0,16507 | 0,43332 | -0,25323 | 0,488862 | -0,15209 | 0,56637 | -0,24679 | 0,2621 | common |
| rs12248176 | chr10:30335464 | 0,115829 | 0,58075 | -0,05112 | 0,914561 | -0,06663 | 0,895356 | 0,1731 | 0,423611 | rare |
| rs2505084 | chr10:30335520 | 0,396745 | 0,17706 | 0,02093 | 0,931401 | 0,307866 | 0,332584 | 0,114771 | 0,566492 | common |

**Data S1. (separate file)**

Supporting material for the results
